# RNase T2-involved selective autophagy of ribosomes induced by starvation in yeast

**DOI:** 10.1101/2023.07.14.548783

**Authors:** Atsushi Minami, Kohei Nishi, Rikusui Yamada, Gai Jinnai, Hikari Shima, Sakiko Oishi, Hirofumi Akagawa, Toshihiro Aono, Makoto Hidaka, Haruhiko Masaki, Tomohisa Kuzuyama, Yoichi Noda, Tetsuhiro Ogawa

**Affiliations:** Department of Biotechnology, The University of Tokyo, Yayoi, Bunkyo-ku, Tokyo 113-8657, Japan; Agro-Biotechnology Research Center (AgTECH), The University of Tokyo, Yayoi, Bunkyo-ku, Tokyo 113-8657, Japan; Collaborative Research Institute for Innovative Microbiology (CRIIM), The University of Tokyo, Yayoi, Bunkyo-ku, Tokyo 113-8657, Japan

**Author notes:** Joint Authors (Atsushi Minami, Kohei Nishi, Rikusui Yamada). Toshihiro Aono, Institute for Agro-Environmental Sciences, National Agriculture and Food Research Organization. 3-1-3 Kannondai, Tsukuba, Ibaraki 305-8604, Japan.

## Abstract

RNase T2 is ubiquitous across diverse organisms, playing essential roles despite its simple enzymatic activity. In *Saccharomyces cerevisiae*, RNase T2, known as Rny1p, localizes in vacuoles and mediates rRNA degradation during autophagy of ribosomes. In this study, we elucidated novel aspects of ribosome degradation mechanisms and the function of Rny1p. First, we discovered that most ribosomes are degraded by selective autophagy, where Rsa1p is the specific receptor of ribosomes to be degraded. Complex structure prediction suggested that Rsa1p also interacts with Atg8p. Furthermore, we observed that the accumulation of rRNA in vacuoles, due to the lack of Rny1p, leads to a decrease in bulk autophagic activity. This decrease in autophagic activity may explain the inability of Rny1p-deficient strains to adapt to starvation conditions. Second, our structural prediction and biochemical analyses indicate that a C-terminal extension, characteristic in fungal RNase T2 including Rny1p, is not necessary for rRNA degradation but for anchoring to the cell wall. Together with molecular phylogenetic analysis, a species-specific role of RNase T2 conferred by the C-terminal extension is suggested.

## INTRODUCTION

Cells comprise numerous proteins; ribosome serves as the platform for synthesizing these proteins. About 2,000 ribosomes are estimated to be produced per minute in cells in the exponential growth phase. Remarkably, the volume occupied by these ribosomes constitute up to 40% of the total cell volume. Since protein synthesis consumes enormous amounts of energy, the cell constantly monitors and controls the activity of ribosomes to balance the energy status. When nutrients are abundant, cells actively synthesize ribosomes. Conversely, when the nutrient source is depleted, ribosome synthesis is promptly inhibited. Simultaneously, the initiation step of protein synthesis is also inhibited. These series of responses prevent energy wastage in a nutritionally restricted environment.

Previously, it was reported that ribosomes are degraded in response to rapamycin treatment in *Saccharomyces cerevisiae* (1). Rapamycin induces a nutrient starvation response in cells by inhibiting TORC1 (<u>T</u>arget <u>O</u>f <u>R</u>apamycin <u>C</u>omplex <u>1</u>), which plays a key role in nutrient source sensing. This indicates that upon nutrient starvation, in addition to suppressing new ribosome synthesis and translation initiation, intracellular ribosomes undergo degradation. However, the details of the degradation mechanism remain unclear.

We have studied the molecular function of the conserved ribonuclease RNase T2. RNase T2 is present in almost all organisms with genomes sequenced, including viruses. RNase T2 is an acidic enzyme with non-specific cleavage activity on single-stranded RNA. Notably, it has been demonstrated that most of RNase T2 degrade rRNA (2–5). Even though the function of RNase T2 is simple, it plays diverse and essential roles in organisms. For example, in humans, RNase T2 has been suggested to function as a cancer suppressor (6). Subsequently, deletions and mutations of the *RNASET2* gene have been reported in patients with cystic leukoencephalopathy (7). Furthermore, RNase T2 is implicated in innate immunity (8–11). In plants, RNase T2, also known as S-RNase, is involved in self-incompatibility, which discriminates between self and non-self cells in Solanaceae, Rosaceae, and Plantaginaceae (12, 13). In viruses, RNase T2 attacks the immune cells of the infected host (14–16). RNase T2 is also present in budding yeast and is called Rny1p. Rny1p possesses a signal peptide and an extension domain, the functions of which remain unknown, at the N- and C-terminal, respectively (17). Additionally, glycosylation sites have already been identified (18) (Figure 1A). This RNase is stress-responsive; for example, Rny1p is required for growth at high temperatures and under high osmotic pressure (17).

**Figure 1.**
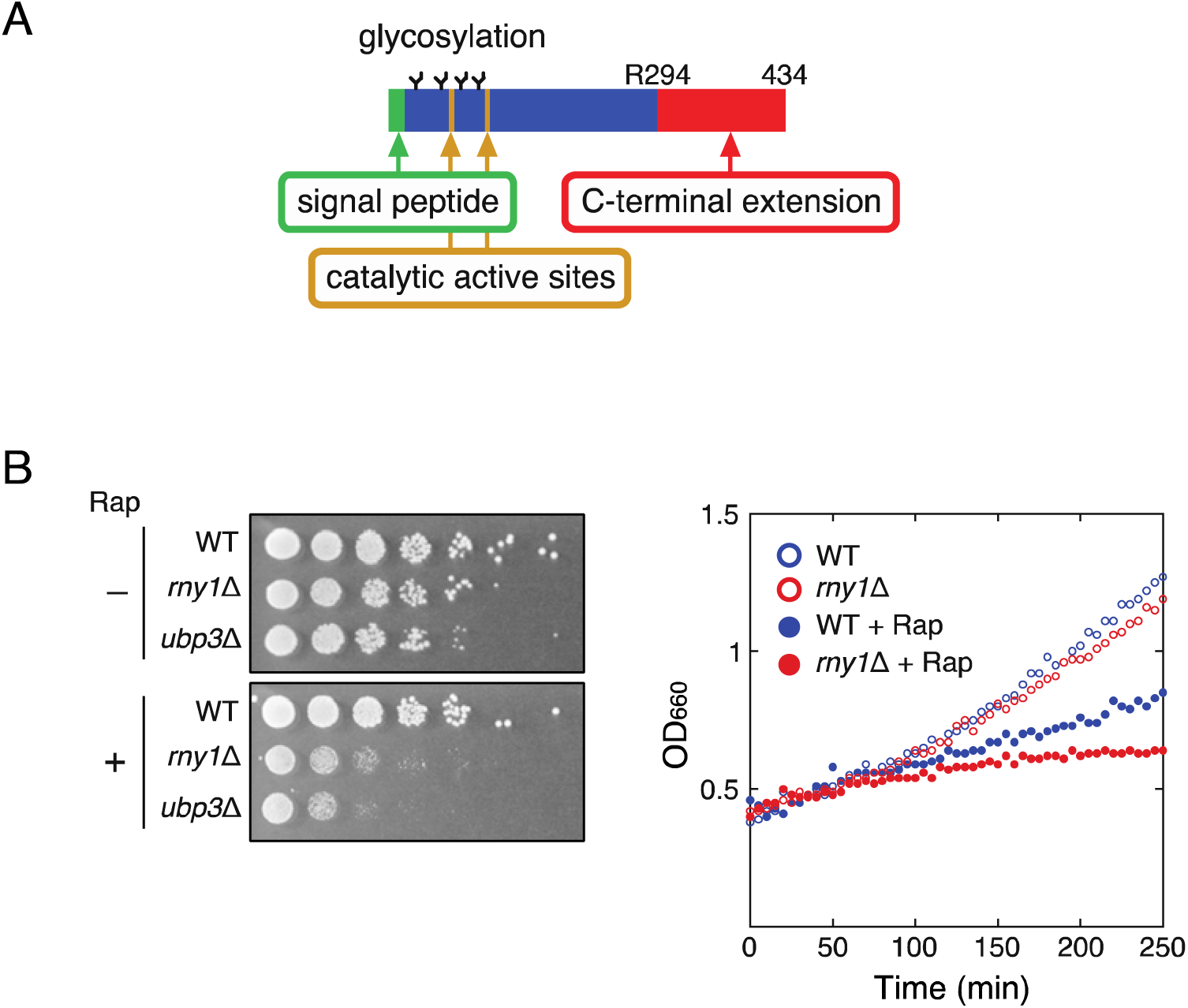
A strain lacking Rny1p is sensitive to rapamycin treatment. (A) The schematic structure of Rny1p is shown. In addition to a signal peptide at the N-terminus, Rny1p possesses a C-terminal extension the function of which is unknown. The glycosylation, catalytic active sites (CASI and CASII), and amino acid numbers from the N-terminus are indicated. CASI and CASII contain catalytic residues His87 and His160, respectively. (B) The wild and *rny1*Δ strains in the log phase were serially diluted, spotted on a solid medium containing 5 nM rapamycin, and incubated at 30 °C. The growth of the *rny1*Δ strain is severely impaired, while the wild-type strain was not affected (left panel). Similarly, growth was measured over time after adding 100 nM rapamycin to wild and *rny1*Δ strains in the log phase. The growth of the wild-type strain was slowed down, while the *rny1*Δ strain stopped growing (right panel). These results showed that Rny1p is required for growth in the presence of rapamycin. Rap means rapamycin to indicate the addition of antibiotics.

Here, in analyzing the function of Rny1p, we present novel insights into ribosomal degradation in response to nutrient starvation mediated by autophagy. Autophagy is a conserved mechanism in which a bilayer membrane, known as autophagosome, envelops the targeted object, transports it to the vacuole, and facilitates its degradation. Two types of autophagy are known: non-selective (bulk) and selective autophagy pathways. The former degrades cytoplasmic components randomly, while the latter selectively recognizes and degrades specific factors. Our results indicate that ribosomal degradation depends mainly on selective autophagy. Selective autophagy requires receptors to specify factors for degradation, and we discovered that Rsa1p, involved in the assembly of 60S ribosomal subunits, functions as the ribosome-specific receptor. We also verified that this selective ribosome degradation differs from the previously reported mechanism known as “ribophagy” (19). Furthermore, we propose a molecular mechanism for ribosome degradation based on biochemical experiments and structural prediction data. In the process, Rny1p deficiency induces rRNA accumulation in the vacuole, which results in the decrease of autophagic activity and failure to adapt to starvation. Finally, we discuss the possibility that the C-terminus of RNase T2 may confer species-specific functions according to the molecular phylogenetic analysis and the biochemical and bioinformatic analysis.

## MATERIALS AND METHODS

### Yeast strains, plasmids, and culture conditions

The strains used in this study are listed in Supplementary Table S1. The pRS316[GFP-ATG8] plasmid, expressing Atg8p fused with GFP at the N-terminus, was provided by the National Bio-Resource Project (NBRP) – Yeast, Japan. The strain chromosomally expressing Rpl25p-GFP (20) was gifted by Professor Hiroshi Takagi. The plasmids pGMH10 and pGMH20 were provided by RIKEN BRC through the National Bio-Resource Project of the MEXT, Japan (cat. RDB01955 and RDB01956, respectively). The strains were cultured in YPD (1% yeast extract, 2% polypeptone, and 2% glucose) or synthetic dextrose (SD) medium (0.67% nitrogen base without amino acids, and 2% glucose) at 30 °C. For gene expression under the control of the *GAL1* promoter, cells were cultivated overnight in a synthetic medium containing raffinose instead of glucose (SRaf medium). After washing, the cells were inoculated into a synthetic galactose (SG) medium to achieve an optical density at 660 nm (OD_660_) of 0.1, and cultivation was re-initiated. Rapamycin (100 nM) was used at the mid-log phase (OD_660_=0.3-0.4) to induce cell starvation. A nitrogen-depleted (SD (-N)) medium containing 0.17% yeast nitrogen base without amino acids and ammonium sulfate, and 2% glucose was used to induce nitrogen starvation.

### Reagents

Rapamycin (a mixture of isomers) was purchased from FUJIFILM Wako Pure Chemical Corporation and dissolved in DMSO. LongLife Zymolyase was purchased from G-Bioscience, and Endoglycosidase H (Endo H) was purchased from New England Biolabs.

### Plasmid construction

Genes for wild-type Rny1p and those lacking the C-terminal extension (Rny1pΔC) without stop codons were PCR amplified, introducing EcoRI and XhoI sites at the 5′ and 3′ ends, respectively. Subsequently, they were digested with EcoRI and XhoI and inserted into the same restriction sites of pYN735, resulting in fusion genes with a 3xFLAG tag coding region at the 3′ ends. These genes were then excised with EcoRI and SalI and inserted into the same restriction sites of pGMH20, a high-copy number plasmid containing the *GAL1* promoter and *HIS3* marker. The resulting plasmids, pGMH20-*RNY1*-*F* and pGMH20-*RNY1*ΔC*-F*, express Rny1p and its variant lacking the C-terminal extension, respectively, each with a 3xFLAG-tag fused at their C-termini. A plasmid expressing catalytically inactive Rny1p was prepared by introducing mutations to exchange the 87th and 160th histidine residues with phenylalanine using the QuickChange site-directed mutagenesis method. The resulting plasmid was designated pGMH20-*RNY1(H87F/H160F)*-*F*. These plasmids were then introduced into BY4742 strains, and the genes were expressed under the control of the *GAL1* promoter.

### Strain construction

Wild-type *RNY1* and *HIS3* cassettes were PCR amplified to introduce homologous regions at the 5′ and 3′ sides of the genomic *RNY1*. These homologous regions were incorporated into the amplified fragment at both ends, using pGMH20-*RNY1-F* as a template. The amplified fragment was transduced into the BY4742 strain, and the genomic *RNY1* gene was replaced by double-crossover homologous recombination. Transformants were plated onto SD medium containing leucine, lysine, and uracil at 30 °C and incubated until visible colonies were obtained. Transformants in which the native *RNY1* gene was successfully replaced with the transduced gene were screened using colony PCR. Similarly, DNA fragments containing genes encoding Rny1pΔC or Rny1p (H87F/H160F) and the *HIS3* cassette were amplified from pGMH20-*RNY1*ΔC*-F* or pGMH20-*RNY1(H87F/H160F)*-*F* as described and integrated into the genome. The *HIS3* cassette was replaced with the *RNY1* gene to construct the *rny1*Δ strain, which serves as a control. The *RNY1* gene of these transformants was then PCR amplified and sequenced to verify the proper integration of the transduced fragments.

### Measurement of cell growth

Cells were grown in a YPD medium at 30 °C until OD_660_ reached 0.3–0.4. These cultures were serially diluted five times, spotted on plates containing rapamycin (5 nM), and cultivated at 30 °C until visible colonies were formed. To assess the effect of starvation on cell growth over time, the wild-type and *rny1*Δ strains were grown in a liquid YPD medium until the OD_660_ reached 0.4. Further, rapamycin (100 nM) was added, and cell growth was automatically monitored using an OD-Monitor & T instrument (TAITEC). In all experiments concerning rapamycin treatment, DMSO was used as the solvent for rapamycin and was added as a mock treatment; we confirmed that the concentration of DMSO used here does not affect the growth of the wild-type strains. To examine the extent of RNA degradation in cells cultured in YPD, the OD_660_ was measured every hour, and cell cultures were diluted with YPD to maintain an OD_660_ between 0.85 and 1.3 (1). As the *CDC48* gene is an essential gene, the CDC48-Tet-off strain purchased from the Yeast Tet-Promoters Hughes Collection (Open Biosystems) was used; expression of the *CDC48* was suppressed by the addition of doxycycline (Figure 3). CDC48 Tet-Off cells at an OD_660_ of 0.1 were cultivated with or without 10 µg/mL of doxycycline in a YPD medium. After 12 and 18 h of cultivation, rapamycin (100 nM) was added, and the culture was continued for 2 h. The cells were subsequently sampled, RNA and vacuoles were stained, and fluorescence microscopic analysis was performed.

### RNA preparation

Total RNA was extracted using the hot phenol method. The collected cells were resuspended in 400 µL of AE buffer (50 mM sodium acetate (pH 5.0) and 10 mM EDTA) and 40 µL of 10% SDS, and briefly mixed at room temperature. Further, 500 µL of AE-buffer-equilibrated phenol (pre-warmed at 65 °C) was immediately added, and the samples were vigorously mixed for 1 min. The samples were incubated at 65 °C for 5 min and mixed for 5 s every 30 s. The samples were then centrifugated, and 300 µL of the resulting supernatant was collected. Subsequently, 400 µL of phenol: chloroform (1:1) and 100 µL of water were added, and the samples were vigorously mixed for 1 min. Three hundred microliters of the supernatant was collected, and 400 µL of chloroform:isoamyl alcohol (24:1) and 100 µL of water were added, and the samples were vigorously mixed for 1 min. After centrifugation, 300 µL of the resulting supernatant was collected, and total RNA was recovered by ethanol precipitation. For electrophoresis, the solution volume of rRNA to be loaded onto a denaturing agarose gel was normalized according to cell numbers (estimated by the OD_660_ value) to compare the amount of rRNA per cell. After electrophoresis, the gel was stained with SYBR Gold (Thermo Fisher Scientific) to visualize the RNA.

### Quantification of RNA by real-time RT-PCR

Total RNA, prepared by the hot-phenol method, was purified using the NucleoSpin RNA Clean-up kit (Takara Bio). Subsequently, 100 ng of each RNA sample was reverse-transcribed with PrimeScript RT Master Mix (Perfect Real Time) (Takara Bio). qPCR was performed using the QuantiTect SYBR Green PCR Kit (QIAGEN). The absolute amount of rRNA obtained was calculated using a calibration curve. Quantified rRNA was normalized to the cell number (estimated using the OD_660_ value).

### Western blotting

Whole proteins were prepared according to a previously described method (21). To detect GFP-Atg8p and Rpl25p-GFP, an anti-GFP polyclonal antibody (Medical & Biological Laboratories) and a rabbit horseradish-peroxidase-linked IgG whole antibody (Sigma-Aldrich) were used as the primary and secondary antibodies, respectively. An anti-DYKDDDDK tag, monoclonal antibody (FUJIFILM Wako Pure Chemical Corporation), anti-mouse IgG, and a sheep horseradish-peroxidase-linked whole antibody (GE Healthcare) were used for the detection of C-terminally FLAG-tagged Rny1p. To detect His-tagged Rsa1p, anti 6x histidine MoAb (9C11) (FUJIFILM Wako Pure Chemical Corporation) was used. An anti-Rpl3p monoclonal antibody was prepared using a hybridoma purchased from the Developmental Studies Hybridoma Bank.

### ALP assay

ALP assay was performed according to a previously described method (22). Strain PhoΔ60 was constructed based on the BY4742 background. Protein concentrations were determined using the Bradford method. α-Naphthyl phosphate disodium salt (Sigma-Aldrich) was used as a substrate, and fluorescence was measured at emission and excitation wavelengths of 360 nm and 465 nm, respectively.

### Endo H treatment, concentration of cell culture medium, and preparation of spheroplasts

After 3 h of nitrogen starvation, cells were collected and suspended in 100 µL of SD (-N) medium and 1 µL of 2-Mercaptoethanol, vortexed for 1 min, and then placed on ice. Further, 15 units of LongLife Zymolyase were added, and the mixture was incubated for 1 h at 37 °C. The spheroplast and cell wall fractions were separated by centrifugation. To remove glycosylation, 500 units of Endo H were added and incubated for 1 h at 37 °C. Cell culture concentration was performed using a VIVA SPIN TURBO 15 device (30,000 MWCO; Sartrius).

### Fluorescence microscopic analysis

FM-4-64 Dye (Thermo Fisher Scientific) and GRG post stain 10000× in water (Biocraft) were used to stain the vacuoles and RNA accumulated in the vacuoles, respectively. An Olympus BX40 microscope with a fluorescence system (Olympus) and an FV1200 laser-scanning microscope (Olympus) were used for imaging.

### Statistical analysis

Statistical analyses were performed using JASP software (23).

### AlphaFold2 modeling and structural alignments

The structural models were predicted using the AlphaFold2 Colab web server (https://colab.research.google.com/github/sokrypton/ColabFold/blob/main/AlphaFold2.ipynb) (24, 25) or its local version (https://github.com/YoshitakaMo/localcolabfold) (24, 26). The reliability of the model was estimated using the pLDDT score and PAE matrix. Structural alignments were performed, and figures were constructed using PyMOL 2.5.0 (Schrodinger). The pLDDT score was mapped onto the structure using the PSICO plugin (https://github.com/YoshitakaMo/pymol-psico).

### Phylogenetic analysis

Fifty-five amino acid sequences of proteins belonging to the RNase T2 superfamily listed in Swiss-Prot were obtained using a UniProt search. Homologous amino acid sequences were removed using CD-HIT (27). The amino acid sequence of *Irpex lacteux* RNase T2 (Irp3, GenBank ID: BAC00516.1) was then added. A phylogenetic tree was constructed using the Graph Splitting method (28).

## RESULTS

### Rapamycin treatment-induced rRNA degradation depends on a different pathway than ribophagy

First, the sensitivity of the *rny1*Δ strain to rapamycin was examined. In both solid and liquid cultures, the growth of the *rny1*Δ strain was inhibited by rapamycin compared to the wild strain, suggesting a link between Rny1p and the starvation response to rapamycin (Figure 1B). Next, we examined whether rRNA degradation induced by rapamycin treatment is dependent on Rny1p. 18S and 25S RNAs prepared from rapamycin-treated *rny1*Δ cells were more abundant compared to those from wild-type cells (Figure 2). These results indicate that Rny1p is involved in the ribosome degradation induced by rapamycin.

**Figure 2.**
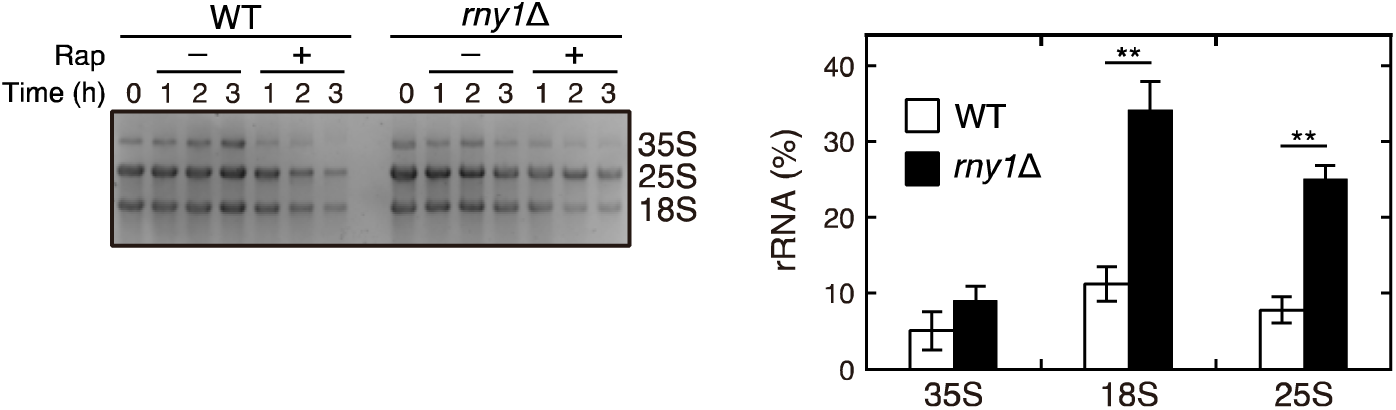
Rny1p degrades 18S and 25S rRNA in response to rapamycin treatment. (A) Wild-type and *rny1*Δ strains were cultivated in duplicate at 30 °C until they entered the log phase. Then rapamycin (100 nM) was added to one of the two cultures, and cultivation was continued. Cells were collected at the indicated time points, and total RNA was extracted. After preparation, total RNA was normalized to the cell number and loaded onto a denaturing agarose gel. (B) Cells were cultivated, rapamycin was added as shown in (A), and further incubated for 3 h. Then total RNA was prepared, and 18S, 25S, and 35S rRNA were quantified by quantitative RT-PCR (qRT-PCR) analysis. The amount of rRNA remaining after rapamycin treatment was calculated as follows: rRNA (%) = 100 × (amount of RNA prepared from rapamycin-treated cells)/(amount of RNA prepared from cells before rapamycin treatment). Data are presented as the means ± standard error of 3 independent experiments. Asterisks indicate significant differences (\*\**P* < 0.01, Student’s t-test). The results in (A) and (B) indicate that 18S and 25S rRNA degradation is Rny1p-dependent. However, the intracellular level of precursor 35S rRNA of the *rny1*Δ strain is almost comparable to that of the wild-type strain, indicating that Rny1p degrades mature rRNAs.

During our study, Huang *et al*. reported the metabolome of RNA degradation caused by nitrogen starvation in detail, particularly from a non-selective autophagy perspective. They also demonstrated that Rny1p is required for the first step of the degradation process (29). We performed a nucleic acid staining of rapamycin-treated wild-type and *rny1*Δ strains as reported (29). Then nucleic acid-derived fluorescence was observed in vacuoles of the *rny1*Δ strain, as observed with nitrogen starvation (Supplementary Figure S1). Previously, ribosomes have also been shown to be degraded by a selective autophagy process known as “ribophagy” in *Saccharomyces cerevisiae* (19). However, we verified that factors required for ribophagy, namely Ubp3p, Ufd3p, and Cdc48p (19, 30), are all dispensable, indicating that the RNA degradation shown here depends on a different pathway than the known ribophagy (Figure 3). Notably, ribophagy experiment also employs nitrogen starvation conditions. Additionally, nitrogen starvation also inhibits TORC1 activity, as does rapamycin. Therefore, the experiments hereafter were conducted under nitrogen starvation conditions instead of rapamycin treatment.

**Figure 3.**
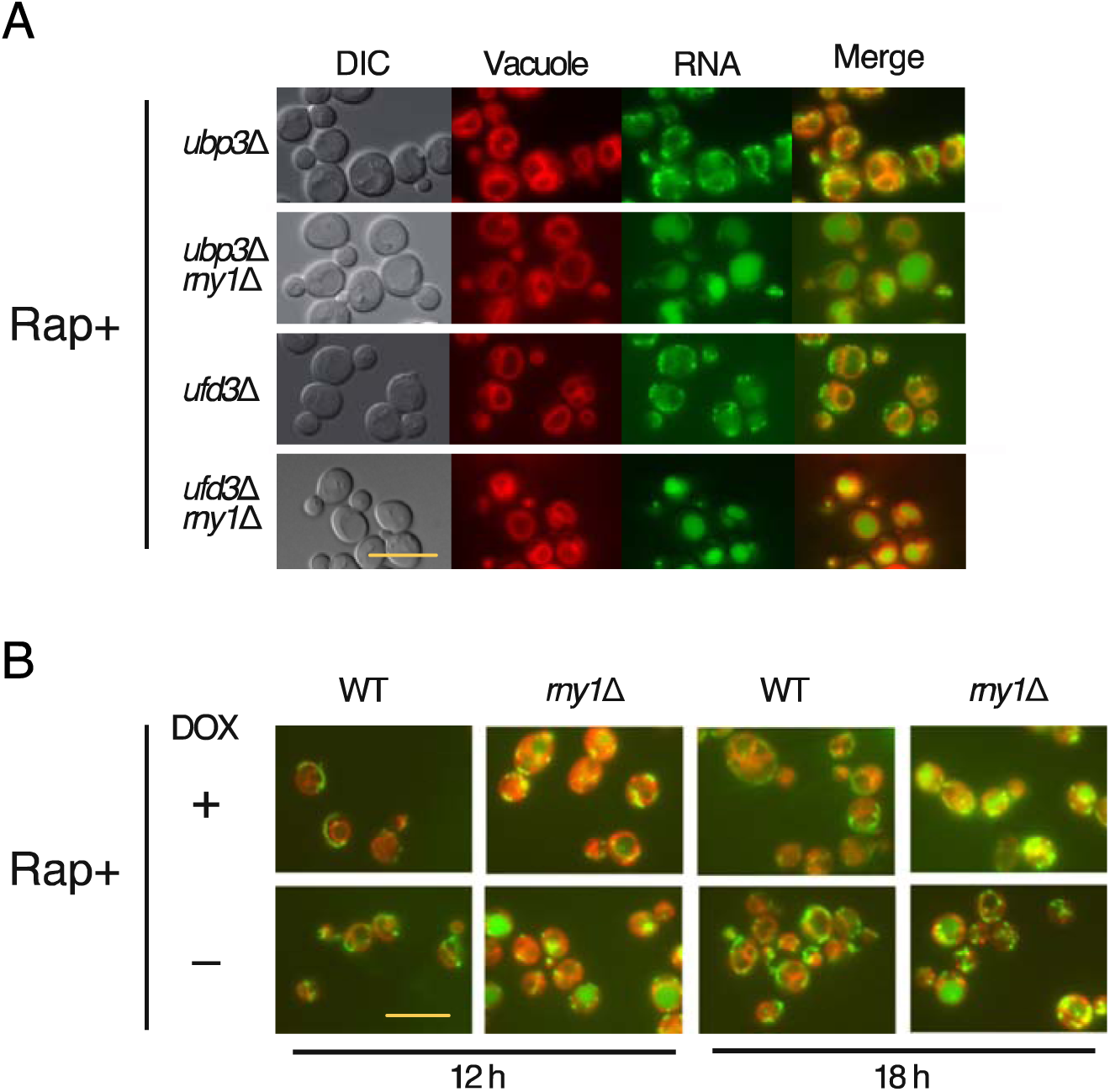
Selective autophagy of ribosomes shown in this study differs from previously reported ribophagy. (A) The strains of *ubp3*Δ, *ubp3*Δ*rny1*Δ, *ufd3*Δ, and *ufd3*Δ*rny1*Δ were grown to the log phase. The cells were cultured with or without rapamycin (100 nM) for 2 h. The vacuoles and RNA were then stained according to Materials and Methods. Further, fluorescence microscopic analysis was performed. Strains *ubp3*Δ*rny1*Δ and *ufd3*Δ*rny1*Δ still accumulate RNA in the vacuole after the rapamycin treatment. (B) CDC48 Tet-Off cells were cultivated in a YPD medium with or without doxycycline. Rapamycin was added at the indicated time, and cells were grown for 2 h. After cultivation, the cells were sampled, and RNA and vacuoles were stained and observed as in (A). RNA fluorescence was observed in the vacuole of CDC48 Tet-Off cells cultivated in the medium containing doxycycline; in the cells, the expression of *CDC48* is suppressed. Together, these results in (A) and (B) indicate that Ubp3p, Ufd3p, and Cdc48p are not involved in the RNA degradation shown in Figure 2. The scale bars in (A) and (B) represent 10 µm.

### Rsa1p involvement in selective ribosome degradation

Huang *et al*. reported that a certain number of ribosomes are non-selectively delivered to the vacuole by autophagy under nitrogen starvation (29). However, as mentioned above, ribosomes are selectively degraded by ribophagy in the same condition (19). Therefore, it remains unclear whether ribosomal degradation is selective, non-selective, or both types coexist. Recently, it was reported that when mTORC1, a homolog of yeast TORC1, is inactivated, ribosomes bind to nuclear fragile X mental retardation–interacting protein 1 (NUFIP1) and are delivered to autophagosomes for selective degradation in mammalian cells (31). We assumed that ribosomes are selectively degraded in yeast and Rsa1p, a functional homolog of NUFIP1 in *S. cerevisiae*, is involved in this process. Rsa1p serves as a platform for box C/D small nucleolar ribonucleoprotein (snoRNP) assembly (32–34) as well as NUFIP1 and localizes to the nucleus.

First, to determine whether the degradation of ribosomal proteins depends on Rsa1p, we induced nitrogen starvation in cells expressing ribosomal protein L25 fused with GFP (Rpl25p-GFP) from the chromosome (20) to perform a green fluorescent protein (GFP) cleavage assay. As autophagy progresses, Rpl25p-GFP is incorporated into the vacuole, resulting in the degradation of Rpl25p. Meanwhile, as GFP is stable in the vacuole, the progress of ribosomal protein degradation can be evaluated by detecting GFP fragments (Figure 4A). The number of GFP fragments increased in wild-type cells over time. Contrarily, the fragment was barely detectable in *rsa1*Δ cells, with a faint band observed after 6 h of nitrogen starvation. Further, we investigated the involvement of Rsa1p in rRNA degradation using fluorescence microscopy (29) (Figure 4B). In *rsa1*Δ cells, RNA accumulation in the vacuoles was not observed. After 3 h of nitrogen starvation, RNA accumulation was slightly observed in *rsa1*Δ*rny1*Δ cells. We quantified RNA degradation in each cell type using qRT-PCR (Figure 4C). Although *RNY1* was intact, RNA was more abundant after starvation in *rsa1*Δ cells than in wild-type cells. The results in Figures 4A, 4B, and 4C demonstrate that the degradation of both ribosomal protein and rRNA degradation is Rsa1p-dependent and selective. Moreover, when Figure 3 is considered together, a new type of selective autophagy on ribosome degradation is suggested.

**Figure 4.**
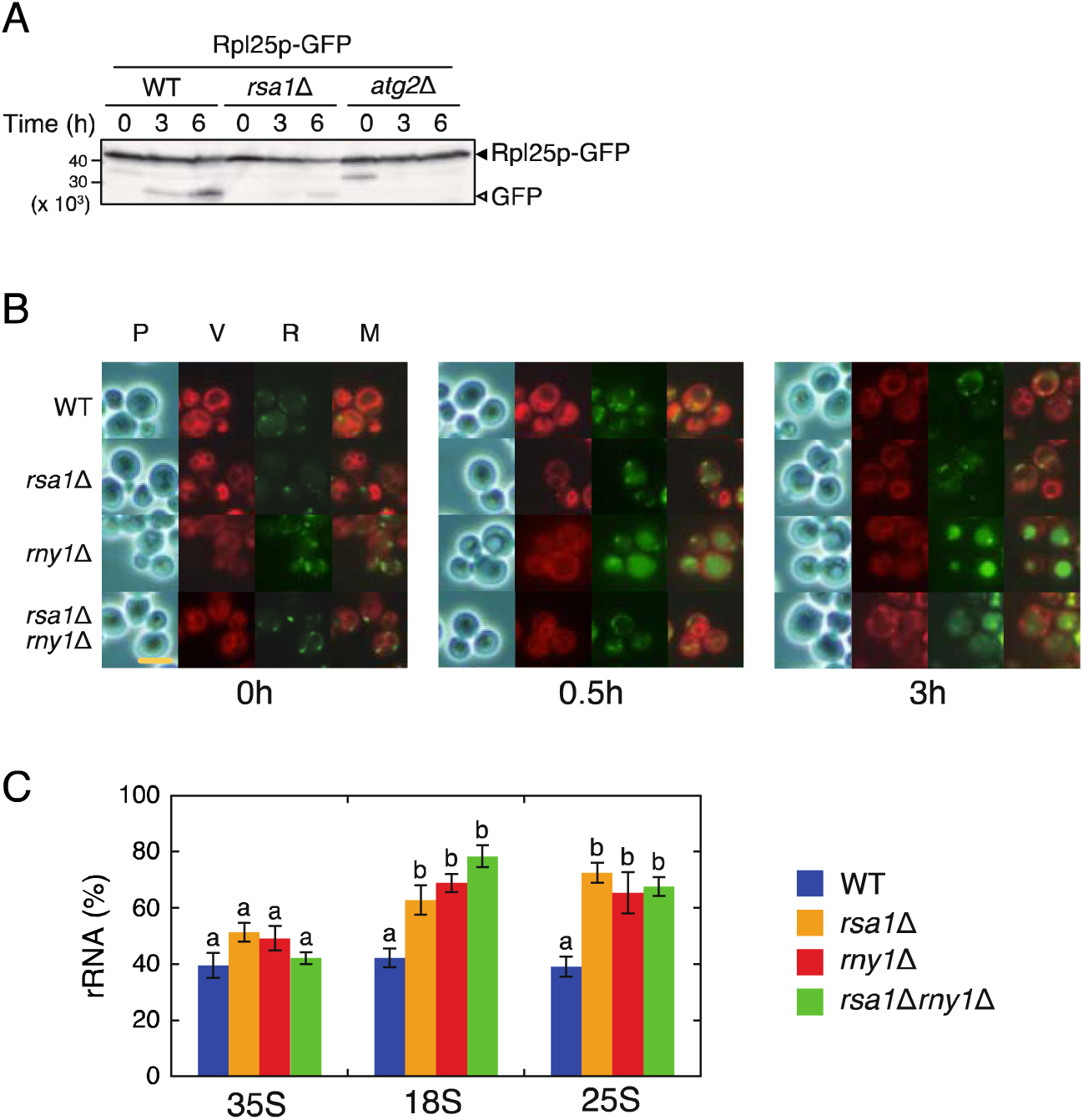
Most ribosome degradation induced by nitrogen starvation depends on Rsa1p. (A) Wild-type, *rsa1*Δ, and *atg2*Δ strains expressing the GFP-tagged ribosomal protein (Rpl25p-GFP) from a chromosome were cultured to the log phase. Then nitrogen starvation was induced, and cell lysates were prepared at the indicated time points. Western blotting was performed using an anti-GFP antibody, and Rpl25p degradation was evaluated. Bands of GFP fragments were observed depending on the incubation time in the wild-type strain. On the other hand, in the *rsa1*Δ strain, a faint GFP band was detected 6 h after starvation induction. This result indicates that Rpl25p degradation is Rsa1p dependent. (B) Wild-type, *rsa1*Δ, *rny1*Δ, and *rsa1*Δ*rny1*Δ strains were grown under nitrogen starvation conditions, and vacuoles and RNA were stained at the indicated time points. “P” labeled above the panels indicates the observation was made by phase-constant microscopy, while “V” and “R” indicate the observation of cells in which vacuoles or RNA were stained, respectively. “M” indicates a merged panel with “V” and “R.” In the *rsa1*Δ strain, as in the wild strain, no RNA fluorescence was observed in the vacuole even 3 h after induction of nitrogen starvation. Simultaneously, RNA fluorescence was faintly observed in the *rsa1*Δ*rny1*Δ strain. The scale bar represents 5 µm. (C) Wild-type, *rsa1*Δ, *rny1*Δ, and *rsa1*Δ*rny1*Δ strains were grown in SD or nitrogen-depleted SD medium (SD (-N) medium). After 3 h of cultivation, total RNA was prepared, and amounts of 35S, 25S, and 18S rRNA were quantified by qRT-PCR analysis, as shown in Figure 2. The amount of rRNA remaining after induction of nitrogen starvation was calculated as follows: rRNA (%) = 100 × (amount of RNA prepared from cells cultured in SD (-N) medium)/(amount of RNA prepared from cells cultured in SD medium). Data are presented as means ± standard error from 4 independent experiments. Statistical analyses were conducted for 35S, 18S, and 25S rRNA. Different letters indicate significant differences (*P* < 0.05, Tukey-Kramer test). Compared to wild-type strains, 18S and 25S rRNA levels were higher in the *rsa1*Δ, *rny1*Δ, and *rsa1*Δ*rny1*Δ strains. However, the amount of the precursor 35S rRNA was not significantly different among all strains. The results in (B) and (C) indicate that Rsa1p is involved in RNA degradation, and mature RNA is not a target of Rsa1p. These results shown in Figures 4A, B, and C indicate that Rsa1p is involved in the starvation-induced degradation of both ribosomal protein and rRNA.

### The structural alignment model and the predicted structure supports the proposed role of Rsa1p

If Rsa1p is a ribosome receptor for selective autophagy, the ribosome should have a binding region with Rsa1p. It has been suggested that NUFIP1 interacts with the 60S subunit of ribosomes. However, the specific binding sites remain unidentified (31), and the homology between NUFIP1 and Rsa1p is low. Therefore, we analyzed the Rsa1p binding site of ribosomes by structural alignment based on known structures. Previously, the crystal structure of the substructure of Rsa1p (Rsa1p_234-290_) complexed with Snu13p has been determined (PDB ID: 4NUT) (35). Snu13p is a conserved RNA-binding protein involved in mRNA splicing and rRNA maturation. This protein is a member of the L7Ae superfamily, and the ribosomal proteins eL8A and eL8B in *S. cerevisiae* are also included in the family. eL8A and eL8B are paralogous proteins; however, in the crystal or Cryo-EM structures of yeast ribosomes deposited in the Protein Data Bank (PDB), all instances of eL8 are registered as eL8A. Moreover, the AlphaFold2 predicted structure of eL8B is almost identical to eL8A. Therefore, eL8A is hereafter used and described as Rpl8p to conform to the notation of other ribosomal proteins. The structure of Rpl8p from the Cryo-EM structure of *S. cerevisiae* 80S ribosome (PDB ID: 6GQV) aligns well with that of Snu13p (2.7Å of the root mean square deviation (RMSD)) (Figure 5A). Based on this structural alignment, we constructed the model structure of Rsa1p bound to the ribosome. In the model, the interaction site of Rsa1p is located near the surface of the ribosome. Therefore, Rsa1p is inferred to bind to the 60S subunit of the ribosomes through Rpl8p.

**Figure 5.**
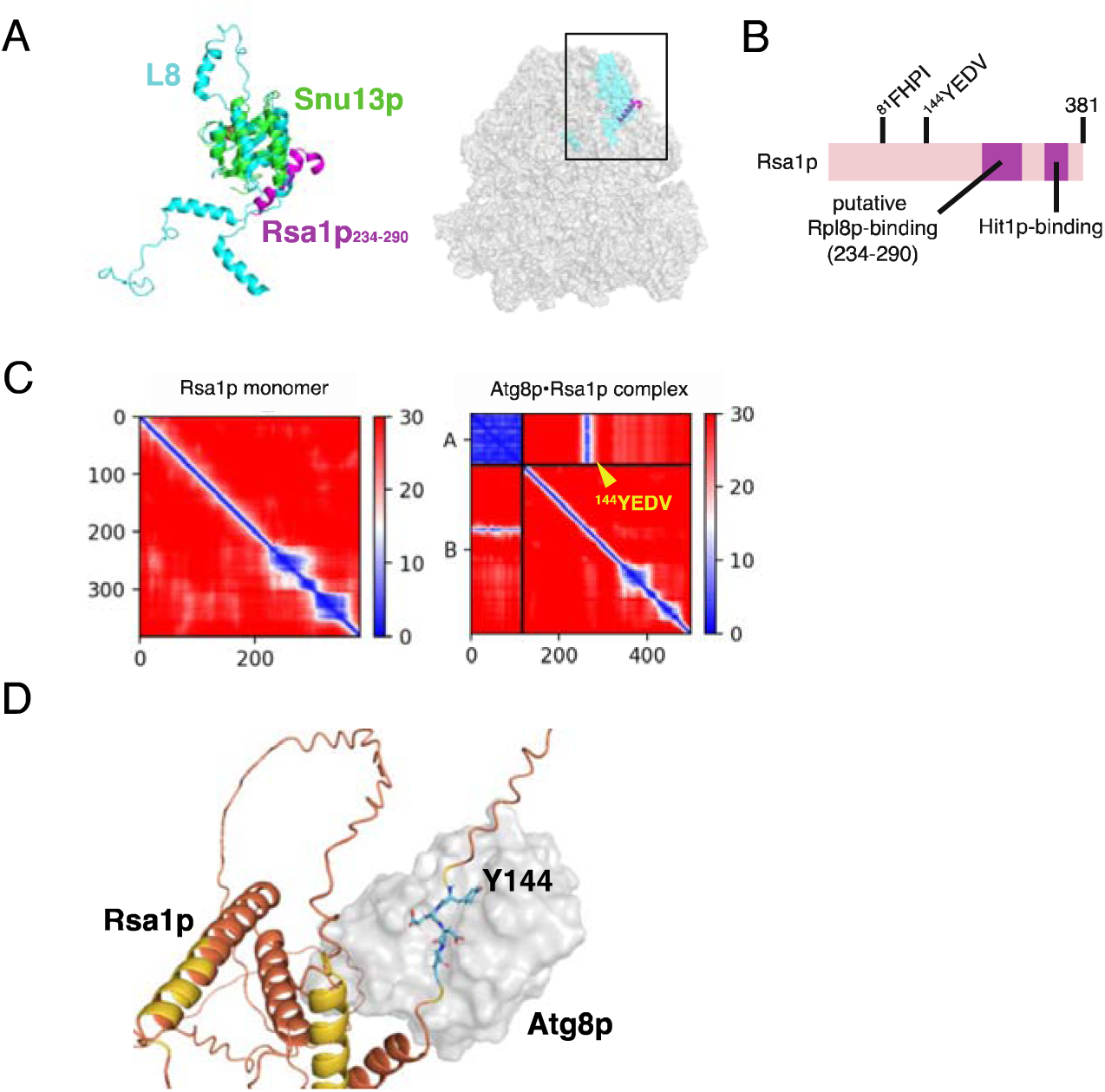
The structural prediction substantiates the Rsa1p-dependent selective ribosomal degradation pathway. (A) Structural model of the complex structure of Rsa1p and the ribosome. In the left panel, the structural alignment of the ribosomal protein L8 and the crystal complex structure of Snu13p and Rsa1p is shown. The Rsa1p-bound *S. cerevisiae* 80S ribosome model is constructed from this alignment and displayed in the right panel. The square in the figure indicates the assumed contact site predicted from the alignment shown in the left panel. (B) Schematic depicting the localization of the putative Atg8p-binding residues in Rsa1p. (C) The PAE matrixes of the AlphaFold2 prediction of Rsa1p in isolation (left panel) and the Rsa1p and Atg8p (right panel) complex. The yellow triangle indicates the putative Atg8p-binding residues in Rsa1p. (D) The predicted complex structure of Rsa1p (*ribbon* with confidence levels mapped onto the structure) and Atg8p (*surface*, gray).

A receptor for selective autophagy is required to bind Atg8p, facilitating the engulfment of cargo by the autophagosome. Atg8p, a ubiquitin-like protein, is essential for autophagosome membrane formation and selective recognition of degradation substrates. In mammalian cells, NUFIP1 was demonstrated to interact with LC3B, a homolog of *S. cerevisiae* Atg8p, upon starvation (31). The LC3-interacting region (LIR) motif, which is also known as the Atg8p-interacting motif represented by [W/F/Y]xx[L/I/V] (36), is essential for this binding of NUFIP1 to LC3B. As mentioned, the sequence similarity of NUFIP1 and Rsa1p is low, and the residues crucial for the binding with LC3B in NUFIP1 are not conserved in Rsa1p (Supplementary Figure S2). Instead, two LIR motifs, ^81^FHPI and ^144^YEDV, were found in Rsa1p (Figure 5B). Accordingly, we tried to predict the complex structure of Rsa1p and Atg8p using AlphaFold2 (24, 37). When the structure of Rsa1p alone was predicted, most regions were disordered, and the total Predicted aligned error (PAE) was relatively low. However, the PAE of residues within ^144^YEDV, one of the candidates for the Atg8p-binding region in Rsa1p, significantly improved in the prediction structure of Rsa1p complexed with Atg8p (Figure 5C). Moreover, the ^144^YEDV residues were located near the cavity of Atg8p in the predicted complex structure, suggesting that ^144^YEDV is a binding site to Atg8p (Figure 5D).

### Impairment of rRNA degradation decreases bulk autophagic activity

We tried to elucidate the involvement of rRNA degradation by Rny1p in the stress response. It was reported that the degradation products of ribosomes are not recycled (37), indicating that rRNA degradation is not aimed at obtaining a nutrient source from the degradation products. In *Arabidopsis*, RNase T2 (RNS2) depletion increases autophagosome, suggesting the activation of autophagy to facilitate the rRNA accumulation in the vacuole (38). We assumed that autophagy activation would also occur in the *rny1*Δ strain; however, it failed as the vacuole was occupied by rRNA, which decreased autophagic activity in *rny1*Δ cells. Autophagic activity can be evaluated by examining the degree of degradation of Atg8p. Therefore, we performed a GFP cleavage assay to measure autophagic activity, as shown in Figure 4A. GFP-Atg8p was expressed in wild-type, *rny1*Δ, and *atg2*Δ strains, and nitrogen starvation was induced. Almost all the GFP-Atg8p was cleaved to GFP fragments after 6 h of starvation in the wild-type strain. Simultaneously, approximately half of the GFP-Atg8p remained intact in the *rny1*Δ strain (Figure 6A). Moreover, prolonged nitrogen starvation increased GFP-Atg8p levels in *rny1*Δ cells, as confirmed by fluorescence microscopic analysis (Supplementary Figure S3). However, the fluorescence intensity of GFP in the vacuole appeared to be lower in many *rny1*Δ cells compared to wild-type cells (Supplementary Figure S3), suggesting that autophagosomes do not fuse efficiently with vacuoles in *rny1*Δ cells. To quantify the autophagic activity of wild-type and *rny1*Δ cells under nitrogen starvation in detail, we performed the alkaline phosphatase (ALP) activity assay. The ALP assay utilizes a strain expressing Pho8Δ60p, a cytoplasmic form of the vacuole-localized alkaline phosphatase Pho8p, that becomes activated in the vacuole. Autophagic activity can be evaluated by measuring the activity of Pho8Δ60p, which is incorporated into the vacuole by autophagy. The ALP activity was lower in *rny1*Δ cells than that in wild-type cells (Figure 6B), indicating that the absence of Rny1p-mediated rRNA degradation reduces autophagic activity. The RNA accumulation in vacuoles may explain the reason for the *rny1*Δ strain not adapting to starvation.

**Figure 6.**
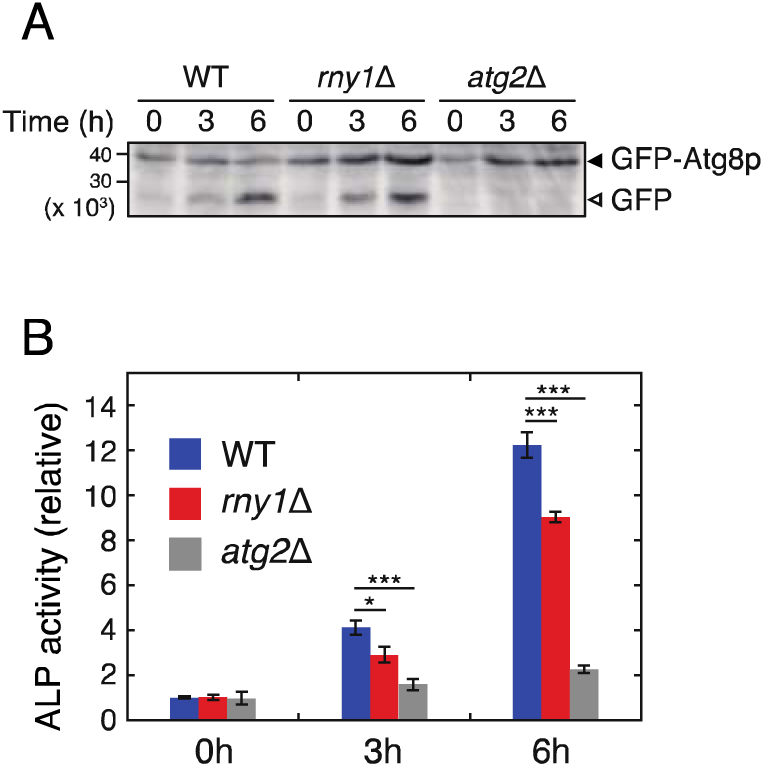
Autophagic activity is decreased in the *rny1*Δ strain. (A) Wild-type, *rny1*Δ, and *atg2*Δ strains expressing GFP-Atg8p were grown in SD medium to log phase. These strains were then cultured in SD (-N) medium and collected at the indicated time points. Cell lysates were prepared, and a GFP cleavage assay was performed. Solid and open arrowheads indicate GFP-Atg8p fusion proteins and cleaved GFP fragments. After 6 h of starvation, most GFP-Atg8p was cleaved to GFP in wild-type strain. However, only half of GFP-Atg8p was cleaved in the *rny1*Δ strain. In addition to this, the total GFP-Atg8p level seemed to be higher in *rny1*Δ compared to wild-type one after starvation. (B) Wild-type, *rny1*Δ, and *atg2*Δ strains expressing the cytosolic form of Pho8p (Pho8Δ60) were cultured in SD (-N) medium, and cell lysates were prepared at the indicated time points. Phosphatase activity was quantified to evaluate the autophagic activity (ALP assay). Data are presented as the means ± standard error of 6 independent experiments. Asterisks indicate significant differences (\**P* < 0.05, \*\*\**P* < 0.001, Student’s t-test).

As mentioned, cells possess an enormous number of ribosomes. Therefore, it was hypothesized that selective autophagy of ribosomes mediated by Rsa1p would compete with non-selective autophagy for autophagy-related factors. However, the GFP cleavage and ALP assays showed that the autophagic activities of wild-type and *rsa1*Δ strains are at similar levels after starvation induction (Supplementary Figure S4). These results indicated that Rsa1p does not perturb non-selective bulk autophagy.

### The C-terminal extension of Rny1p is not required for ribosome degradation but is required for cell wall anchoring

As shown in Figure 1A, Rny1p possesses a C-terminal extension with an unknown function. To investigate whether the C-terminal extension is required for rRNA degradation, we replaced genomic *RNY1* with *RNY1-F*, *RNY1ΔC-F*, or *RNY1(H87F/H160F)*-*F* and induced nitrogen starvation in the resulting strains (please refer “Materials and Methods” section). Vacuolar RNA accumulation was observed in *rny1*Δ and Rny1p (H87F/H160F)-F-expressing strains under a fluorescence microscope (Figure 7A). However, RNA accumulation was not observed in the strain expressing Rny1pΔC-F, as well as in the strain expressing Rny1p-F, indicating that the C-terminal extension is dispensable for RNA degradation in the vacuole.

**Figure 7.**
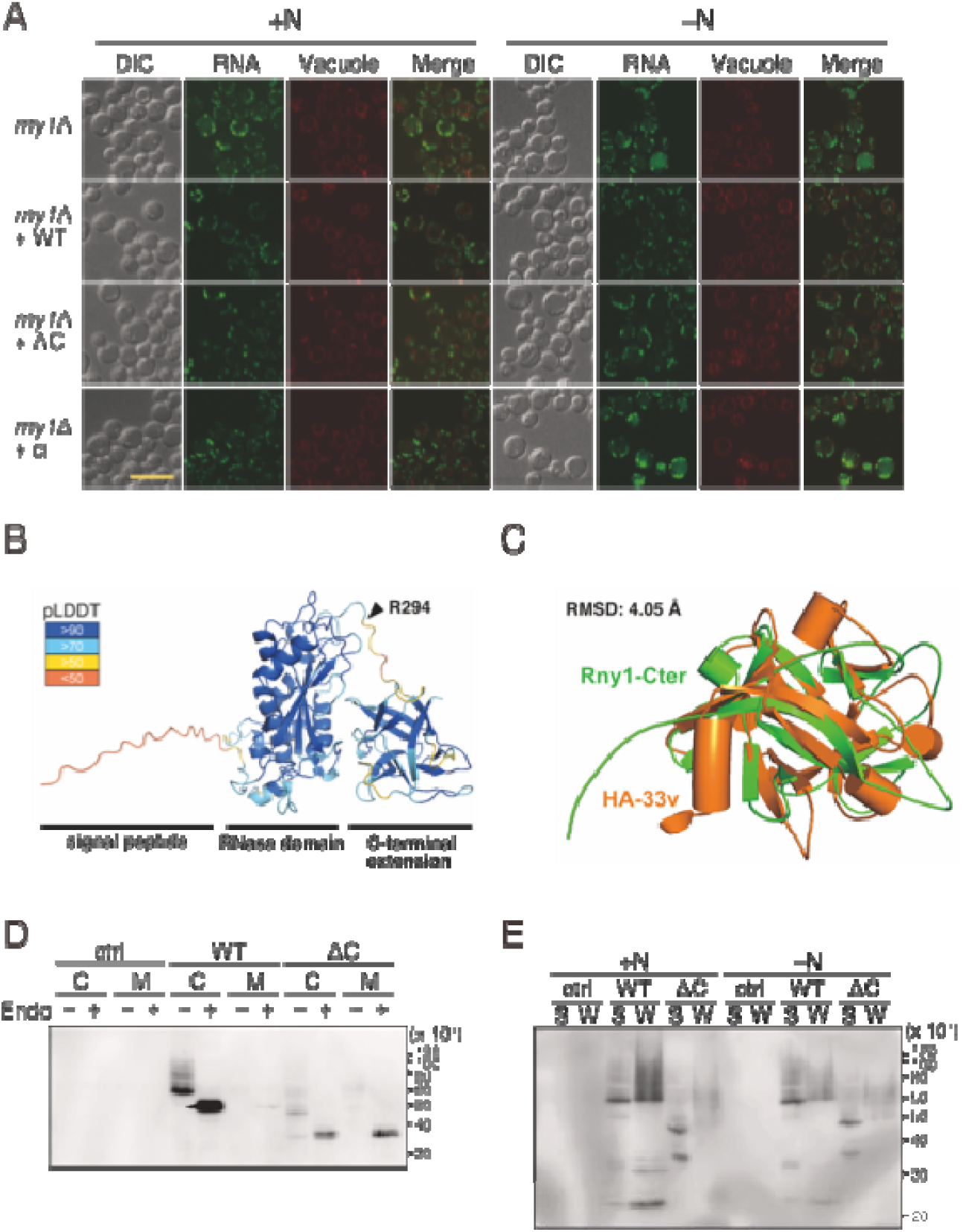
The C-terminal extension of Rny1p is necessary to bind the cell wall. (A) Strains expressing Rny1p-F, Rny1pΔC-F, or Rny1p(H87F/H160F)-F from the chromosomes were cultured in SD or SD (-N) medium for 3 h. After staining the vacuoles and RNA, fluorescence confocal microscopic analysis was performed. RNA accumulation in the vacuole was not observed in cells expressing Rny1p lacking its C-terminal extension, indicating that the extension is dispensable for RNA degradation. The scale bar represents 10 µm. (B) The structure of Rny1p was predicted using AlphaFold2 with confidence levels mapped onto the structure. The signal peptide, RNase domain, and C-terminal extension are indicated. The arrowhead indicates the 294th amino acid residue where the C-terminal extension begins, as shown in Figure 1A. (C) Structural alignment of the C-terminal extension of predicted Rny1p (labeled Rny1-Cter) with the crystal structure of the variable region of HA-33 (PDB ID:5B2H, labeled HA-33v). (D) A Wild-type strain expressing Rny1p-F or Rny1pΔC-F from a plasmid was cultivated in an SG medium. After cultivation, the cells were collected, and cell lysates were prepared. At the same time, the cultured medium was concentrated 200 times by ultrafiltration. Then, western blotting was performed using an anti-FLAG antibody. “C” and “M” indicate cell lysate or concentrated medium loading, respectively, while “Endo” indicates endoglycosidase H treatment. We found that Rny1p lacking its C-terminal extension is secreted to the medium. (E) Strains shown in (D) in the log phase were cultivated with (+N) or without nitrogen source (–N) for 3h. Then, cells were collected and treated with zymolyase to remove the sugar chain of the cell wall. After centrifugation, spheroplasts were collected, and cell lysates were prepared. The final volume of the cell lysate and supernatant containing the zymolyase-treated cell wall fraction was 500 µL and 100 µL, respectively. An equal volume of samples was loaded onto a gel, and SDS-PAGE was carried out, followed by western blotting as shown in (D). “S” and “W” indicate the cell lysates prepared from spheroplasts or cell wall fractions were loaded, respectively. Nitrogen starvation significantly decreases the cell wall-bound form of Rny1p. In (D) and (E), “ctrl,” “WT,” and “ΔC” indicate that samples were prepared from strains harboring pGMH20 (empty vector), pGMH20-*RNY1*-*F,* and pGMH20-*RNY1*ΔC*-F*, respectively.

We analyzed the function of the C-terminal extension from a structural biology perspective due to the lack of information on homologous proteins or known domain motifs other than RNase T2 from the amino acid sequence. According to the overall structure of Rny1p predicted using AlphaFold2 (38) (Figure 7B), the C-terminal extension of Rny1p was distinguishable from the structure of the RNase domain. Further, we searched for similar protein structures as the predicted structure of the C-terminal extension on the Dali server (39). The top five structures (Table S2) are all known or suggested to contain domains involved in sugar binding. The predicted C-terminal extension structure is superimposed to that of the variable region of HA-33 (PDB ID:5B2H) (40), which has the highest Z-score (Figure 7C), despite the low amino acid sequence similarity between them. Rny1p is a vacuole-localized enzyme (29); however, some are secreted extracellularly and bound to the cell wall (18). The structural similarity of the C-terminal extension of Rny1p to sugar-binding domain-containing proteins such as HA-33 suggested that Rny1p binds to the sugar chain of the cell wall via the C-terminal extension.

Furthermore, we experimentally confirmed whether Rny1p binds to the cell wall through its C-terminal extension. Due to the low intracellular level of Rny1pΔC, we prepared cell lysates from the wild-type strains expressing Rny1p-F or Rny1pΔC-F from the high-copy plasmid. Simultaneously, we concentrated the cultured medium by ultrafiltration. As a result of performing western blotting on the lysate and the concentrated culture medium, smeared bands of Rny1pΔC-F, presumably derived from glycosylation, were observed in the medium fraction (Supplementary Figure S5). Rny1pΔC-F was remarkably detected in the Endo H-treated medium fraction, while Rny1p-F was not detected (Figure 7D). These results demonstrate that Rny1p binds to the cell wall through its C-terminal extension. Additionally, from the result of Endo H treatment to cell lysate, Rny1pΔC-F and Rny1p-F are glycosylated (39). However, expression levels were reduced compared to the wild-type. Subsequently, to examine the difference in expression and localization in the presence and absence of a nitrogen source, we prepared spheroplasts from cells cultivated under each condition to separate the cytosolic and cell wall fractions. Western blotting of these fractions revealed a significant reduction in Rny1p-F levels in the cell wall fraction in response to nitrogen starvation. Conversely, no apparent changes were observed in intracellular Rny1p-F levels (Figure 7E).

### The possibility that the C-terminal region provides diversity in the role of RNase T2

We tried to investigate the conservation of the C-terminal extension among RNase T2 family members. As RNase T2 is characterized by two conserved active sites (CASI and CASII), the overall sequence similarity is relatively low. Due to this limitation, we deemed phylogenetic analysis based on standard multiple alignments unsuitable. We used the Graph Splitting method (40) to reconstruct a phylogenetic tree containing the remote homologs instead of the standard multiple alignments. The phylogenetic analysis showed that fungal RNase T2 diverged from the other RNase T2 proteins early in the phylogenetic tree, suggesting that they evolved independently (Figure 8A). A portion of fungal RNase T2, labeled in yellow in the phylogenetic tree, had a long domain exceeding 100 amino acids at the C-terminus (Supplementary Figure S6A). Among them, RNase T2 in *Irpex lacteus*, referred to as Irp3, was reported to share some critical amino acid residues in the C-terminal extension with Rny1p, despite low overall amino acid sequence similarity, suggesting that the C-terminal extensions of RNase T2 may have a similar role to that of Rny1p (41). The predicted structure of the C-terminal extension of Irp3 (Supplementary Figure S6B) superposed well with that of Rny1p, with 2.91 Å of RMSD, which is lower than that of HA-33 (Figure 8B), like those of other fungal RNase T2 proteins (Supplementary Figure S6C and D).

**Figure 8.**
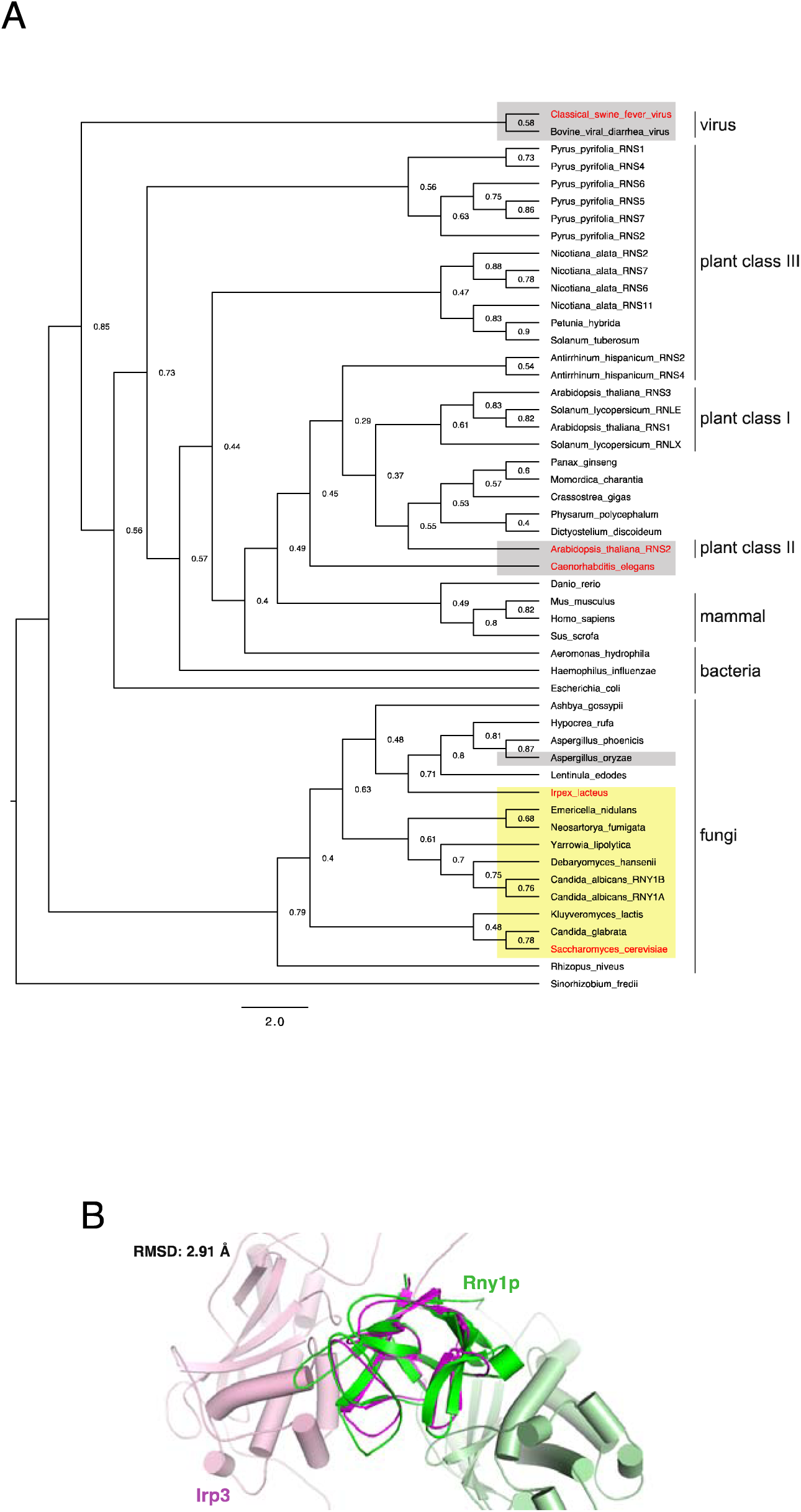
Diversity of the C-terminal extension among the RNase T2 family. (A) A phylogenetic tree of the RNase T2 family was created based on amino acid sequences. Organisms possessing C-terminal extensions homologous to Rny1p are shaded yellow, whereas those not homologous to Rny1p are shaded grey. RNase T2 of classical swine fever (CSF) virus, *Arabidopsis thaliana*, *Caenorhabditis elegans*, *Irpex lacteus*, and *S. cerevisiae* are discussed in this paper, and their genus names are colored in red. *B*, Structural alignment of the C-terminal extension of Rny1p (labeled Rny1p) was performed with that of the predicted *Irpex lacteux* RNase T2 (labeled Irp3).

Contrarily, some RNase T2 proteins, other than those in fungi, also have C-terminal extensions, but their sequences and structures are different from those of Rny1p. For example, E^rns^, a glycoprotein produced by the classical swine fever (CSF) virus, contains a motif characteristic of RNase T2 and exerts RNase activity (14, 42). E^rns^ also has a C-terminal extension that is required for receptor-independent translocation across the cell membrane (43). According to the prediction by AlphaFold2, the C-terminal extension of E^rns^ forms an α-helix structure (Supplementary Figure S6E), which is different from that of Rny1p. Other examples include the C-terminal extension of the class II plant RNase T2, which is necessary for localization to the vacuole (44), and *Caenorhabditis elegans* RNase T2 (45). Their C-terminal extensions are shorter than that of Rny1p and may have evolved independently, based on the phylogenetic tree (Figure 8A). From these analyses, it is inferred that the diversity of the C-terminal extension may result from the evolution of the RNase T2 family, depending on where they function in each organism.

## Discussion

Our findings indicate that most, if not all, ribosomes are selectively degraded in response to nitrogen starvation. This selective ribosomal degradation seems to differ from the process previously reported for ribophagy, as its factors, such as Ubp3, Ufd3p, and Cdc48p (37), are unnecessary (Figure 3). Selective autophagy requires a receptor specific to the degradation target, and our study indicates that Rsa1p plays this role in yeast. Assuming that the amount of degraded rRNA corresponds to that of the degraded ribosome, approximately 30% of ribosomes are estimated to be degraded in a selective autophagy-dependent manner based on the result of Figure 4C. However, bulk autophagy is unaffected by massive ribosomal degradation (Supplementary Figure S4). These results indicate that factors involved in autophagy are present in sufficient amounts in the cell, showing the robustness of the autophagy pathway.

GFP cleavage and ALP assays revealed that the *rny1*Δ strain exhibits a reduction in bulk autophagy activity (Figure 6), indicating that RNA accumulation in vacuoles harms cells, which may result in the growth defect observed in the *rny1*Δ strain under stress conditions. This may be because accumulated RNA inhibits the degradation of other factors that should be degraded in vacuoles. Therefore, in the *rny1*Δ strain, cells may upregulate Atg8p synthesis to facilitate autophagic degradation of RNA (Figure 6 and Supplementary Figure S3) as in *Arabidopsis*. However, autophagosome fusion with vacuoles seems to be reduced in *rny1*Δ cells (Supplementary Figure S3). In *Arabidopsis thaliana*, RNase T2 null mutant accumulates RNA and ribosomes in autophagosomes and autophagic bodies even under a nutrient-rich condition (38). RNA accumulation is also observed in the lysosome of brain neurons in RNase T2-deficient zebrafish (3). The RNase T2 null zebrafish shows an abnormal MRI pattern in the brain, assumed to reproduce the cystic leukoencephalopathy observed in humans (7); this also indicates that the accumulation of RNA in lysosomes is detrimental.

Our discovery of the role of Rsa1p, mediating selective ribosomal degradation, shares similar functions with NUFIP1 in mammals. NUFIP1 forms a complex with the small protein ZNHIT3 (zinc finger HIT domain-containing protein 3) and is involved in the assembly of box C/D snoRNP, as described above. NUFIP1 localizes primarily in the nucleus but is also present in the cytoplasm of certain cell types (47). The inhibition of mTORC1 by starvation or the addition of its inhibitors results in the localization of the NUFIP1-ZNHIT3 complex from the nucleus to autophagosomes and lysosomes. It has been reported that Hit1p, an interacting partner of Rsa1p, is cooperatively involved in box C/D snoRNP assembly (46). Therefore, Hit1p is also assumed to cooperate with Rsa1p to form a receptor for the ribosome in yeast. As mentioned earlier, ribophagy, selective autophagy for ribosome degradation, was first proposed in budding yeast. It involves the ubiquitin protease Ubp3p, Ufd3, and Cdc48, which were shown to be involved in subsequent studies (47). In mammals, the term ribophagy is also used for selective ribosome degradation; however, they did not show whether ubiquitin protease(s), corresponding to yeast Ubp3p, is also required for the NUFIP1-dependent ribosomal degradation (48). Our discovery that Rsa1p mediates Ubp3p-independent selective autophagy of ribosomes suggests that the NUFIP1-dependent ribosomal degradation pathway in mammals may also be distinct from so-called ribophagy.

As described above, new ribosome synthesis and the initiation step of protein synthesis are inhibited when cells are starved. However, some ribosomes should synthesize proteins required for adaptation to stress conditions. Therefore, the random engulfment of cytoplasmic ribosomes by autophagosomes appears to be inefficient. Since NUFIP1 and Rsa1p are involved in ribosome biosynthesis in the nucleus, it is possible that the ribosomes targeted for selective degradation are not those in the cytoplasm but those in the composition process in the nucleus. Under stress conditions, incomplete ribosomes in the nucleus are unnecessary; therefore, they are likely targets of selective degradation. Accordingly, the selective degradation of ribosomes shown here does not interfere with ribosomes attempting to translate stress response factors in the cytoplasm. Alternatively, after translocating from the nucleus to the cytoplasm, Rsa1p may selectively bind to ribosomes involved in translating specific mRNAs, such as those encoding amino acid biosynthesis factors and ribosomal proteins, delivering them to autophagosomes (48). Our structural analysis suggested that ribosomal protein eL8 interacts with Rsa1p. As mentioned, *S. cerevisiae* has two paralogues of eL8; eL8A and eL8B. Although there are few studies on these paralogs, eL8A and eL8B are reported as not functionally interchangeable, and changes in carbon sources alter their relative proportions within ribosomes (49). Therefore, it is presumed that eL8A and eL8B are used differently to bring about different ribosomal functions in response to environmental changes, and Rsa1p might selectively interact with either of them.

Recent advances in protein structure prediction, represented by AlphaFold2, provide new insights in molecular biology, and their applications continue to expand. In particular, applying AlphaFold2 to complexes consisting of multiple interacting proteins, like Rsa1p and Atg8p in this study, seems to be the famous “hack” that was not initially envisioned (50). In this usage, the reliability of its prediction is usually evaluated by the PAE between interacting proteins. For example, in a large-scale screening of protein-protein interactions performed in yeast using complex structure prediction, protein combinations with low PAEs were eliminated from the candidate (51). Rsa1p was considered inherently unsuitable for prediction in this study because of its long-disordered region. The protein complex prediction succeeded, even for the protein with low total PAE. PAEs of specific residues (^144^YEDV, an LIR motif) in Rsa1p showed a distinct improvement in predicting the complex with Atg8p (Figure 5C). To the best of our knowledge, a few applications are present to infer interacting residues from improved PAEs at the level of specific residues within a disordered region rather than at the domain level. This will expand the potential applications of the structural prediction of protein complexes.

After glycosylation, Rny1p is transported to the vacuole, whereas some Rny1p is secreted out of the cell and anchored to the host cell wall (18, 29). Although the C-terminal extension of Rny1p has no homologous sequence other than RNase T2, its structure resembles that of a sugar-binding domain (Figure 7C), suggesting that Rny1p anchors to the sugar chain of the cell wall via the C-terminal extension. The C-terminal extension of Rny1p is unnecessary for RNA degradation (Figure 7A), and we successfully revealed that cell wall anchoring of Rny1p requires the C-terminal extension by biochemical analyses (Figure 7D and Supplementary Figure S5). However, the mechanism by which Rny1p is differentially transported into the vacuole and extracellular space remains unknown. After nitrogen starvation was induced, the amount of extracellular Rny1p decreased, suggesting that starvation changes the cellular trafficking of Rny1p (Figure 7D). The physiological role of cell wall anchoring is also unknown. It is supposed that some extracellular RNAs bind to cell membranes to regulate their ion permeability (52), and Rny1p degrades these RNAs for regulation (17). Our studies have shown that the C-terminal extension is required for growth, at least in high temperatures (Supplementary Figure S7). Anyway, further studies are necessary to clarify the function of the C-terminal extension of Rny1p.

The C-terminal extensions of fungal RNase T2 proteins, including Irp3 produced by *Irpex lacteus*, were predicted to have almost the same structure as Rny1p (Figure 8B). Conversely, the C-terminal extension of E^rns^, which is required to cross cellular membranes in a receptor-independent manner, had a different structure (Supplementary Figure S6E). The Graph Splitting method suggested that fungal RNase T2 evolved independently from animal and plant RNase T2 (Figure 8A). Contrarily, it is noteworthy that the X-ray crystal structures of plant *Pyrus pyrifolia* and *Homo sapiens* RNase T2 were well superposed on the predicted structure of the RNase domain of Rny1p (Supplementary Figure S8). The presence of RNase T2 in almost all organisms suggests that cells have retained RNase T2 since the early stage of evolution. Over its long evolutionary history, RNase T2 has played a specific role in many species. Acquisition of the C-terminal extension may be one of the reasons for diverse roles of enzymatically simple ribonucleases in different species. To better understand this process, the structure and function of the C-terminal extension of the RNase T2 family must be studied.

## ACKNOWLEDGEMENTS

We acknowledge Dr. Hiroshi Takagi (Nara Institute of Science and Technology) for providing the strain expressing Rpl25p-GFP. We thank the National Bio-Resource Project (NBRP) of the MEXT/AMED, Japan, for providing plasmid pRS316[GFP-ATG8]. We also thank RIKEN BRC through the National BioResource Project of MEXT, Japan, for providing the plasmids pGMH10 (RDB01955) and pGMH20 (RDB01956).

---

**Supplementary Figure S1.**
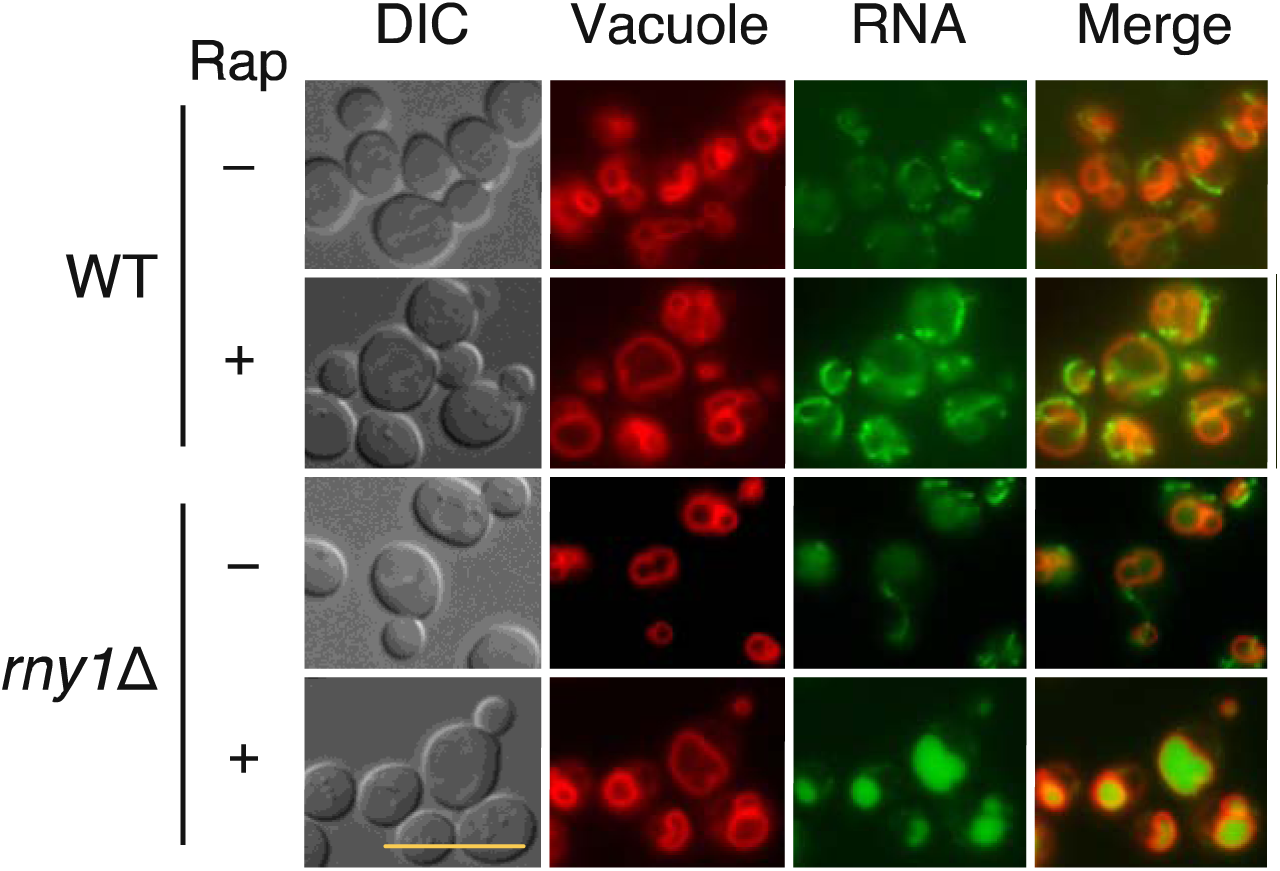
Rny1p degrades RNA in the vacuole upon rapamycin treatment, as well as nitrogen starvation. Cultivation, rapamycin treatment, and cell staining were performed, as in Figure 3. The scale bar represents 10 μm.

**Supplementary Figure S2.**
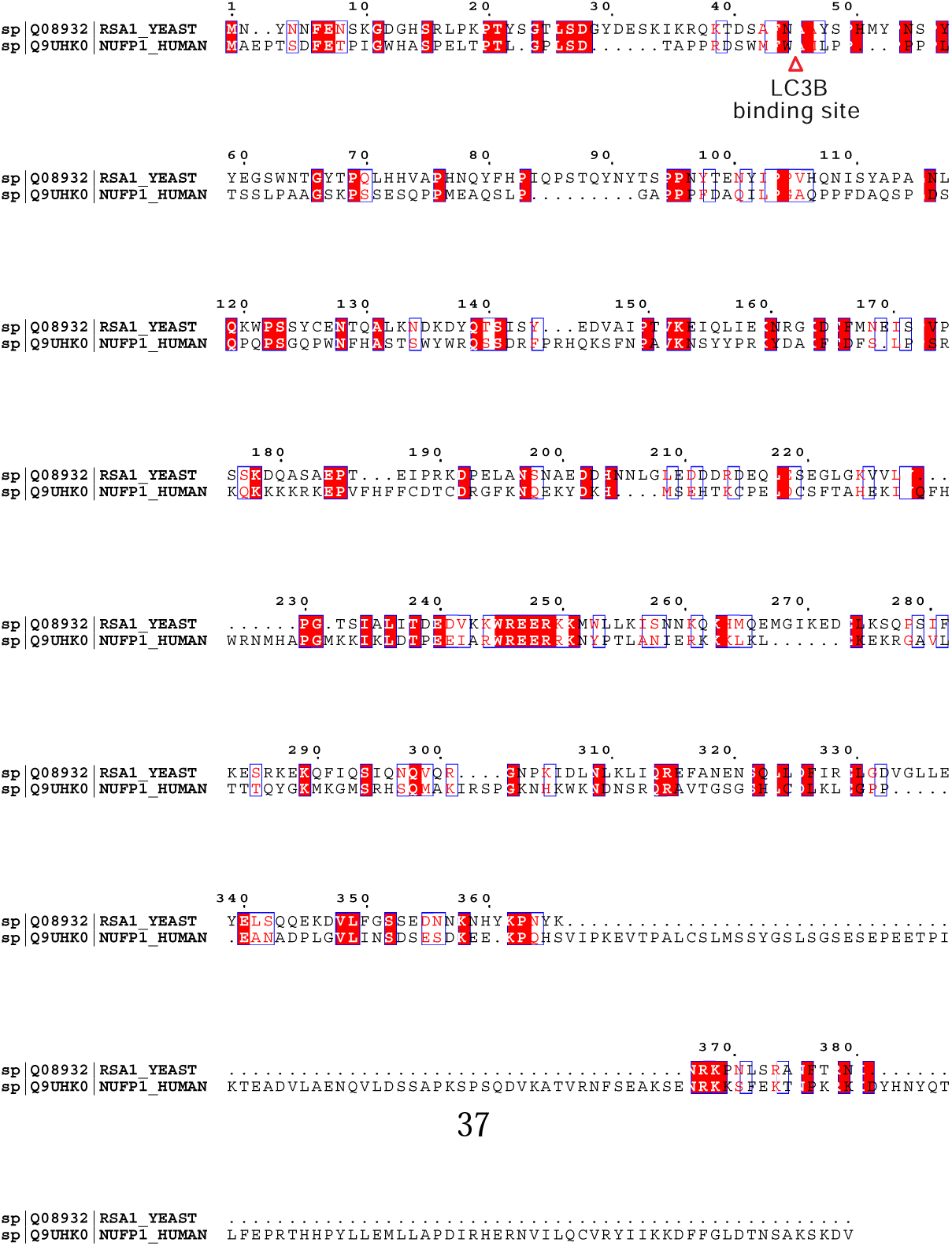
The pairwise sequence alignment of Rsa1p and NUFIP1. The amino acid sequence of Rsa1p is aligned with that of NUFIP1. The LC3B binding site of NUFIP1 is shown in yellow in the figure. W40 was reported to be the residue responsible for binding to LC3B, but this residue is not conserved in the sequence of Rsa1p.

**Supplementary Figure S3.**
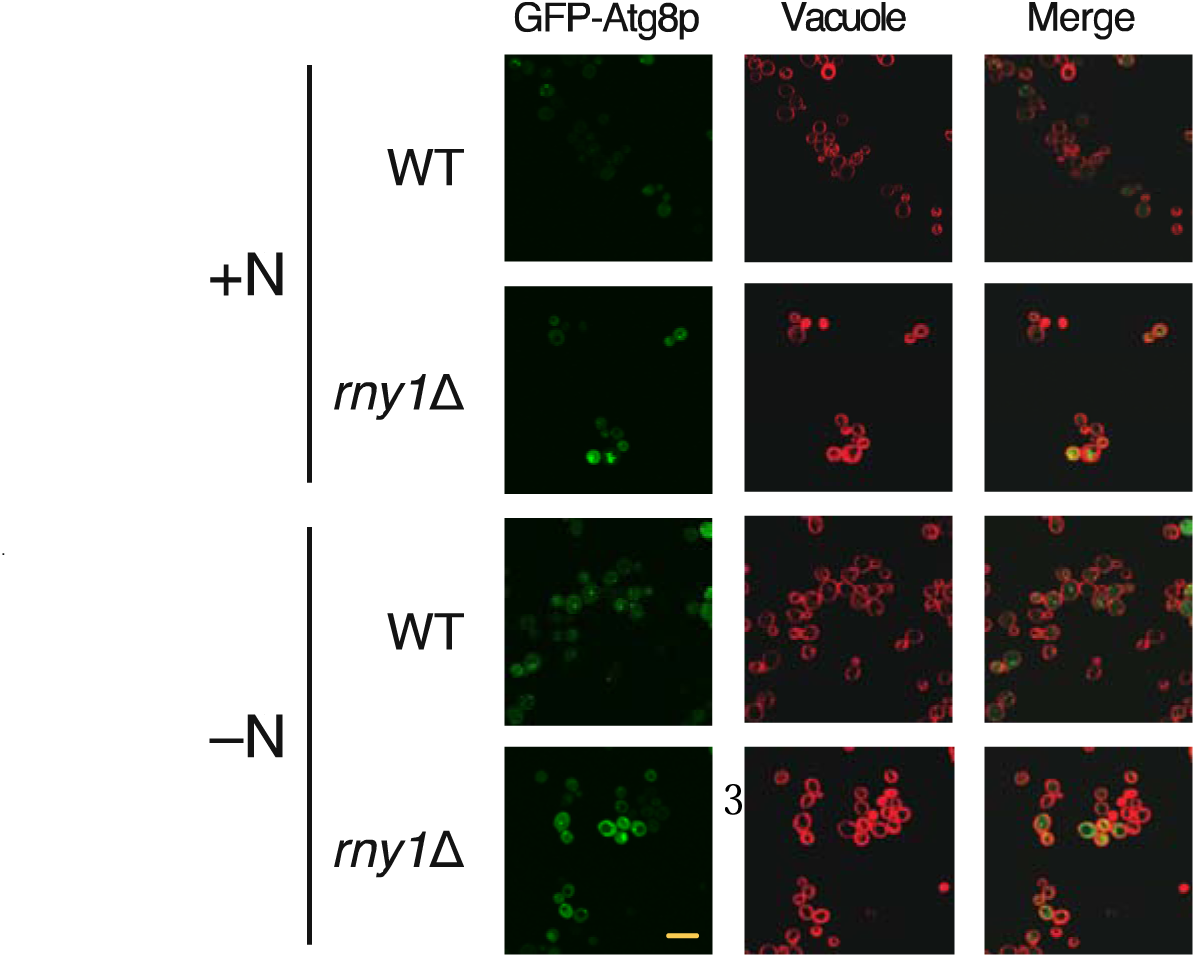
Accumulation of GFP-Atg8p in the vacuole is reduced upon nitrogen starvation in *rny1*Δ cells. Wild-type and *rny1*Δ strains expressing GFP-Atg8p (see “Materials and Methods” section in the main text) were grown in SD medium to the log phase. Further, cells were suspended in SD or SD (-N) medium and cultured for 3 h. After vacuoles were stained, fluorescence microscopic analysis was performed to observe GFP-Atg8p and vacuoles. The scale bar represents 10 μm.

**Supplementary Figure S4.**
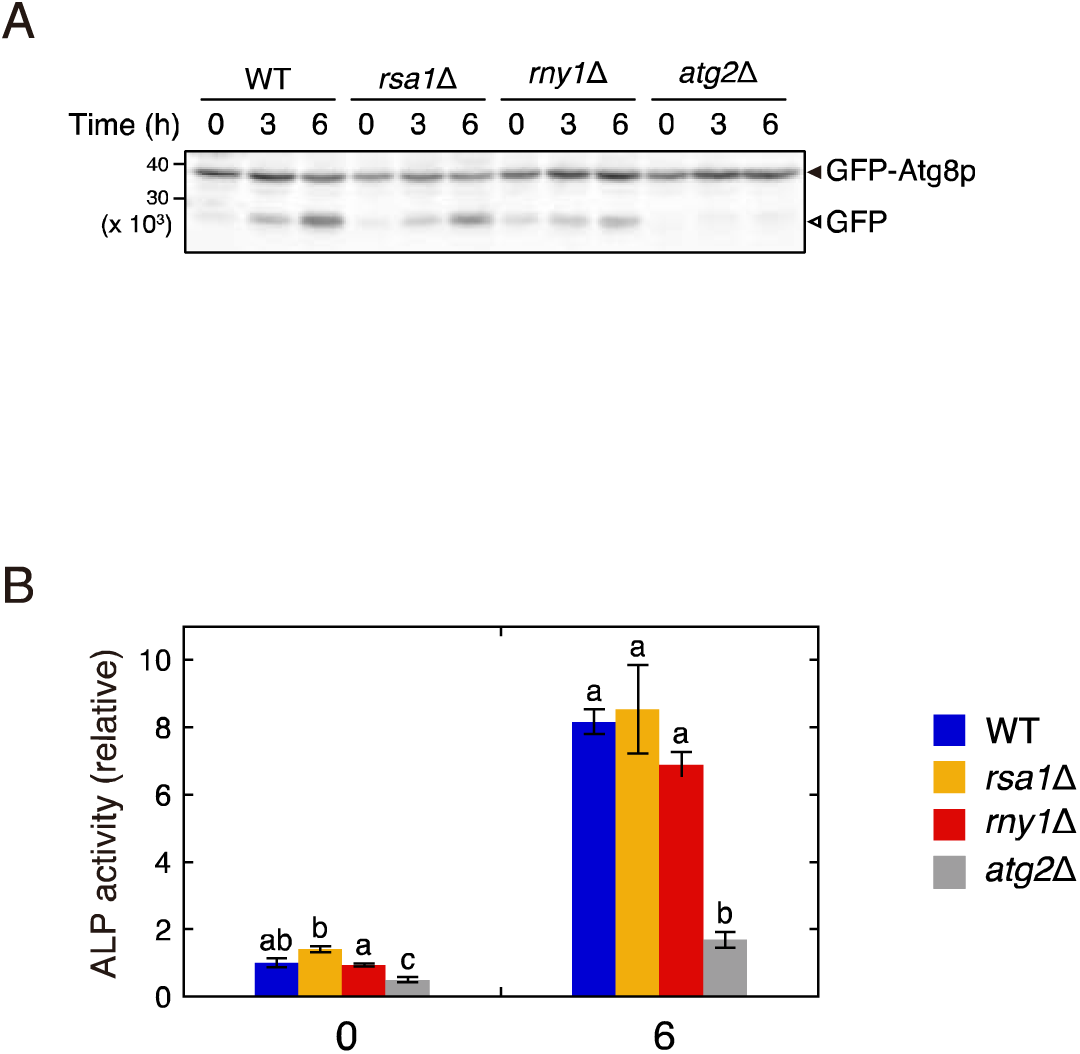
Bulk autophagic activity is not perturbed by Rsa1p-dependent selective autophagy of ribosomes. (A) Nitrogen starvation was induced in wild-type, *rsa1*Δ, *rny1*Δ, and *atg2*Δ strains, as shown in Figure 4, and then the cells were collected at the indicated time points. Cell lysate preparation and western blotting were performed as per the “Materials and Methods” section. GFP-Atg8p and GFP fragments are indicated by closed and opened arrowheads, respectively. No significant differences in the appearance of GFP fragments over time were observed in both wild-type and *rsa1*Δ strains. (B) Nitrogen starvation was induced in wild-type, *rsa1*Δ, *rny1*Δ, and *rsa1*Δ*rny1*Δ strains expressing Pho8Δ60 for 6 h, and then ALP assay was performed. Data are presented as means ± standard error from 3 independent experiments. Statistical analyses were conducted at each point in time. Different letters indicate significant differences (*P* < 0.05, Tukey-Kramer test). The bulk autophagy activity of the *rsa1*Δ strain is almost the same as that of the wild-type strain, indicating that selective autophagy of ribosomes does not interfere with bulk autophagy.

**Supplementary Figure S5.**
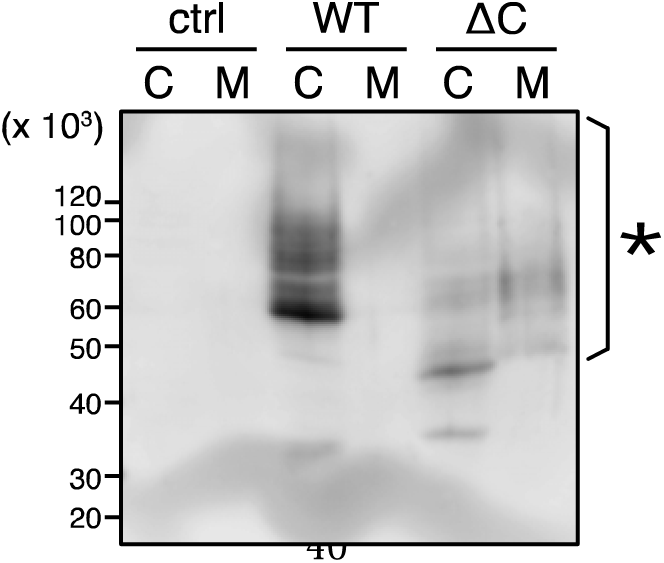
Rny1pΔC-F is glycosylated and released into the medium. The wild-type strain expressing Rny1p-F or Rny1pΔC-F from a plasmid was cultivated. Then, cell lysate and concentrated medium were prepared, and western blotting was performed using an anti-FLAG antibody. The asterisk indicates glycosylated Rny1pΔC-F observed in the concentrated medium. “C” and “M” above the panel indicate that the cell lysate or concentrated medium was loaded, respectively, while “ctrl” indicates that wild-type strain carrying pGMH20 was used as a control.

**Supplementary Figure S6.**
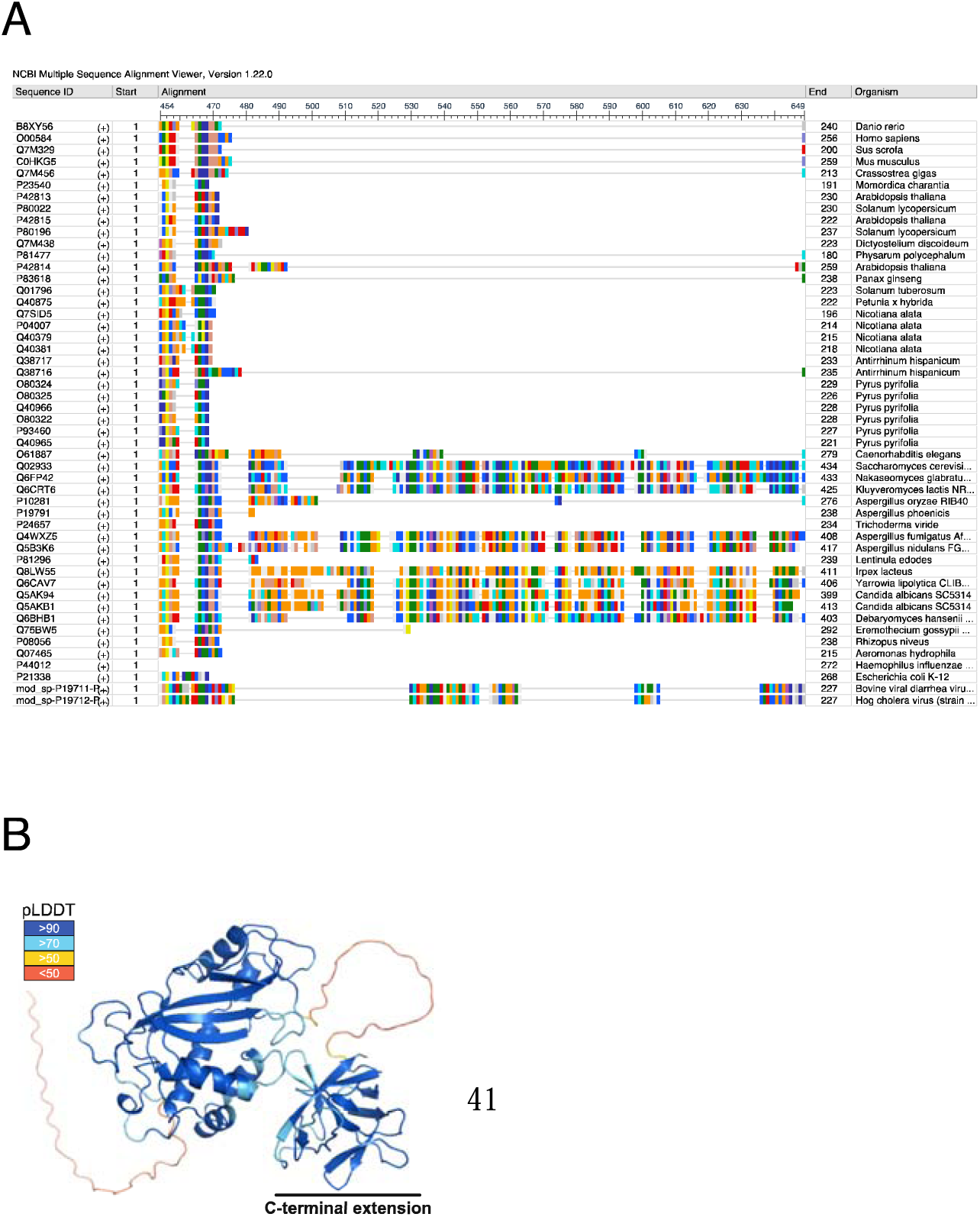
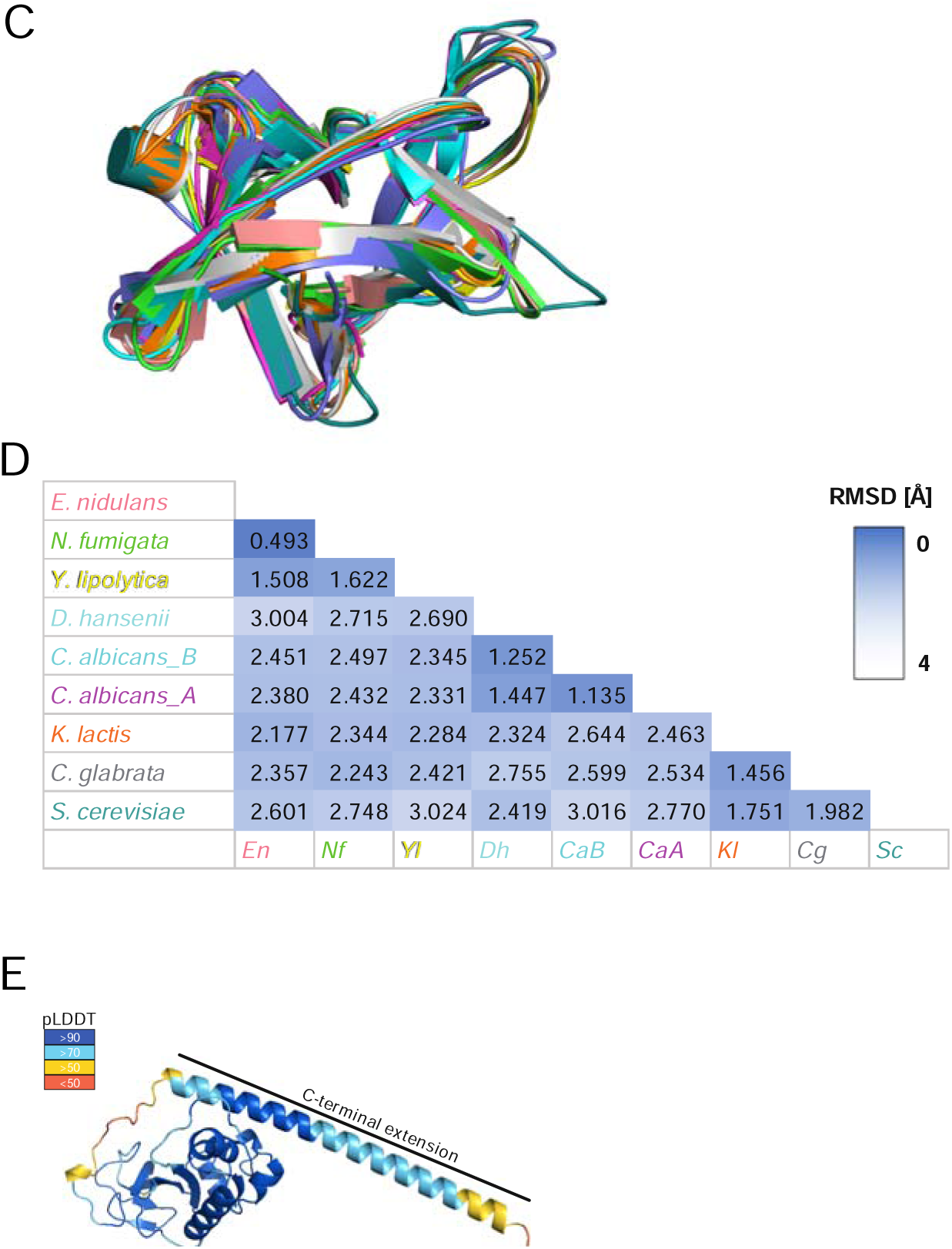
Diversity of the C-terminal extensions of RNase T2. (A) Multiple sequence alignment (MSA) of the C-terminus of the RNase T2 family. MSA was constructed using MAFFT (v7.511, online version) and illustrated using the NCBI MSA viewer (v1.22.0). In the C-terminal extension, each color indicates an individual amino acid. (B) The structure of *Irpex lacteux* RNase T2 (Irp3) was predicted by AlphaFold2, with confidence levels mapped onto the structure. (C) Structural alignment of the predicted structures of the fungal RNase T2 C-terminal extension. Salmon: *Emericella nidulans*; green: *Neosartorya fumigata*; yellow: *Yarrowia lipolytica*; slate: *Debaryomyces hansenii*; cyan: *Candida albicans* RNY1B; magenta: *Candida albicans* RNY1A; orange: *Kluyveromyces lactis*; grey: *Candida glabrata*; deep teal: *Saccharomyces cerevisiae*. (D) RMSD values of (C) for structural alignments. (E) The structure of E^rns^, a CSF virus RNase T2, was predicted using AlphaFold2, with confidence levels mapped onto the structure.

**Supplementary Figure S7.**
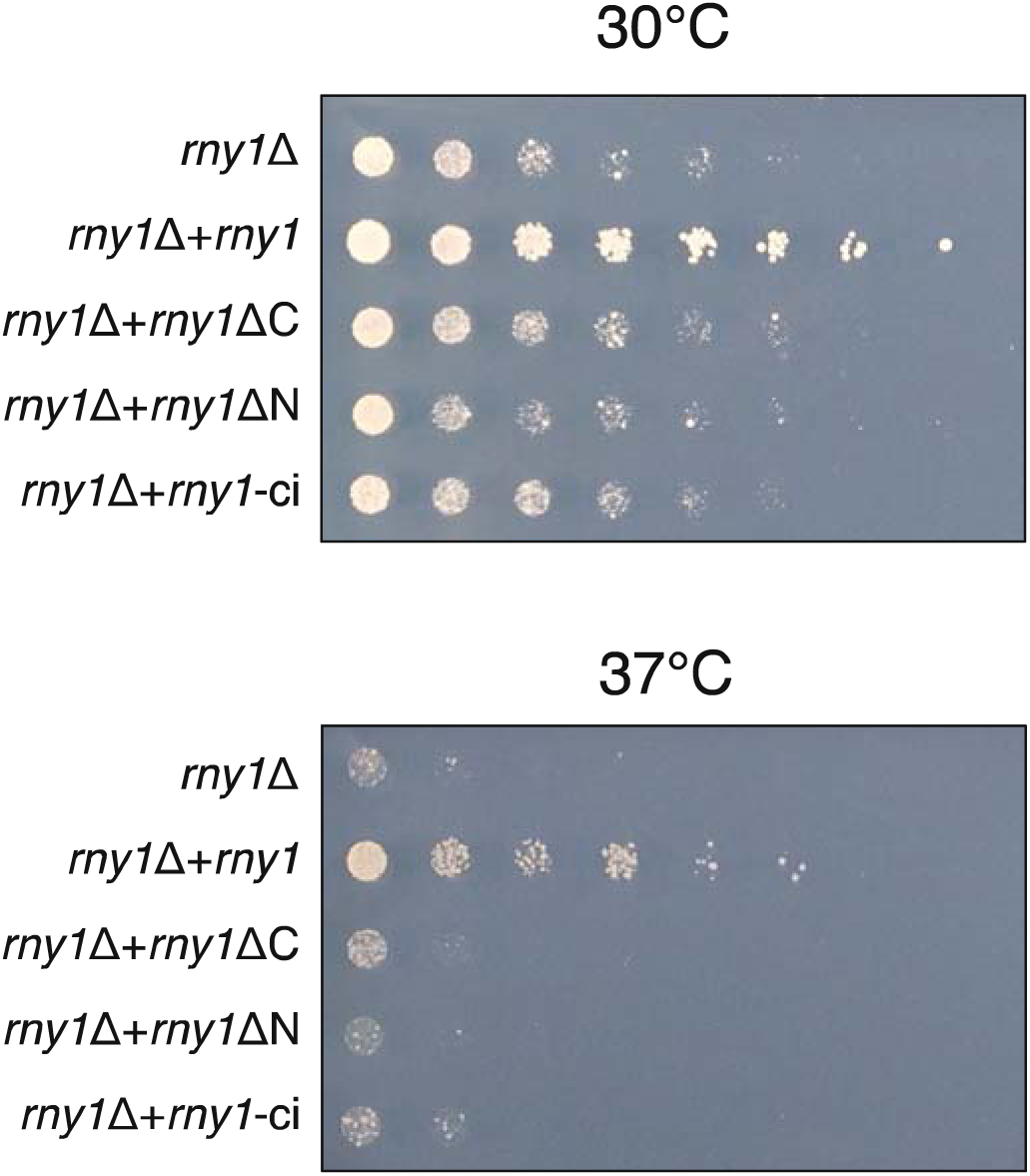
The C-terminal extension is required for growth at high temperatures. *rny1*Δ strains carrying the plasmid pGMH20 encoding Rny1p-F, Rny1pΔC-F, Rny1pΔN-F (Rny1p-F lacking the N-terminal signal peptide), and catalytically inactive Rny1p-F (rny1-ci) were cultured in SG medium to log phase. Further, they were serially diluted 3 times and spotted onto a solid YPD medium. The plates were incubated at 30 or 37 °C.

**Supplementary Figure S8.**
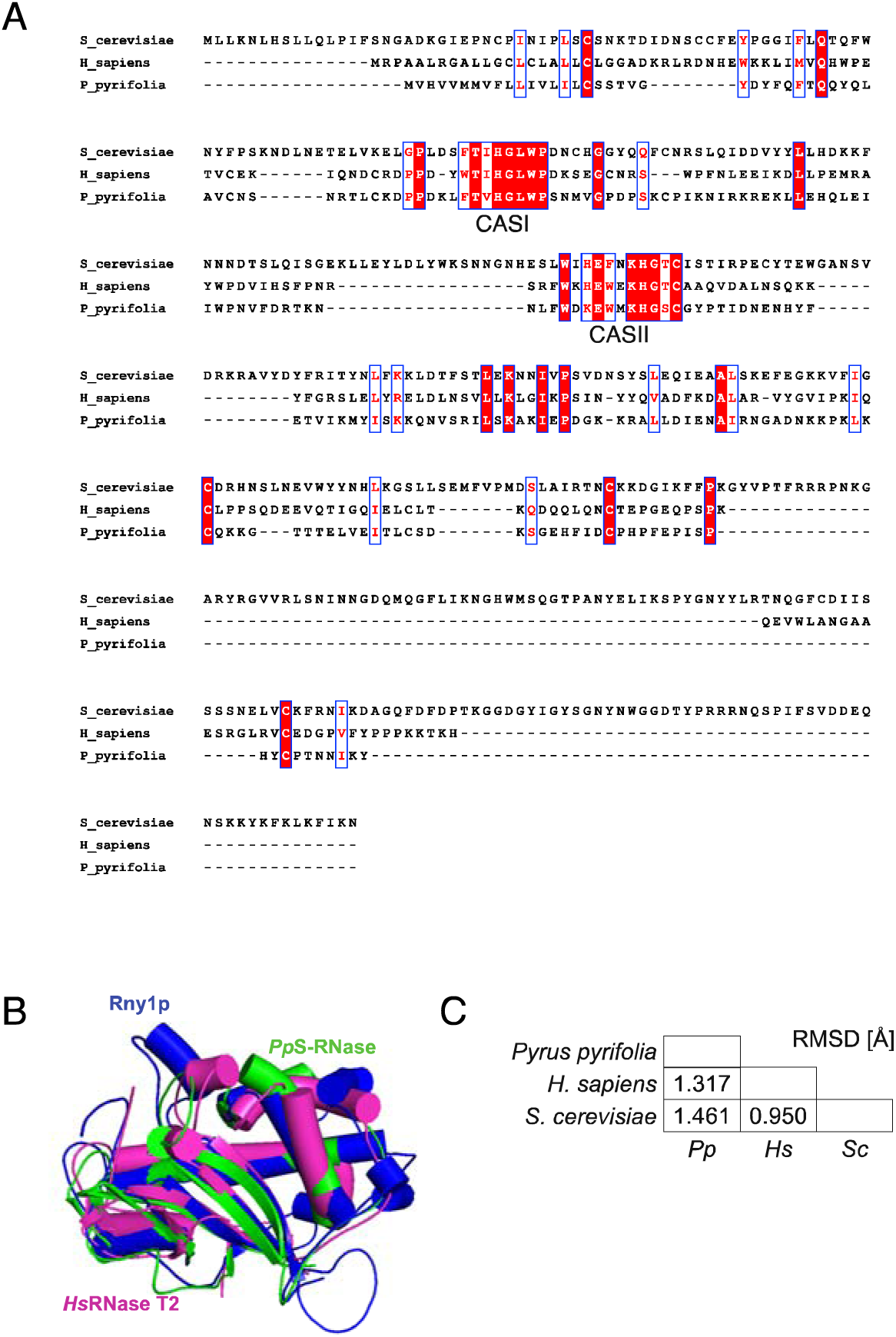
Structural conservation of the RNase domain of eukaryotic RNase T2. (A) MSA of representative RNase T2 sequences in eukaryotes. The sequence was constructed using ClustalW (https://www.genome.jp/tools-bin/clustalw). The labels shown on the left of the amino acid sequences are as follows: S_cerevisiae: RNase domain of Rny1p from *Saccharomyces cerevisiae* predicted by AlphaFold2, H_sapiens: Ribonuclease T2 from *Homo sapiens* (PDB ID: 3T0O), and P_pyrifolia: Ribonuclease S- 3 from *Pyrus pyrifolia* (PDB ID: 1IQQ). (B) Structural alignment of the predicted RNase domain of Rny1p (labeled Rny1p) with the crystal structures of *Homo sapiens* (labeled HsRNase T2) and *Pyrus pyrifolia* (labeled PpS-RNase). (C) RMSD values of (B) for structural alignments.

**Table S1.**
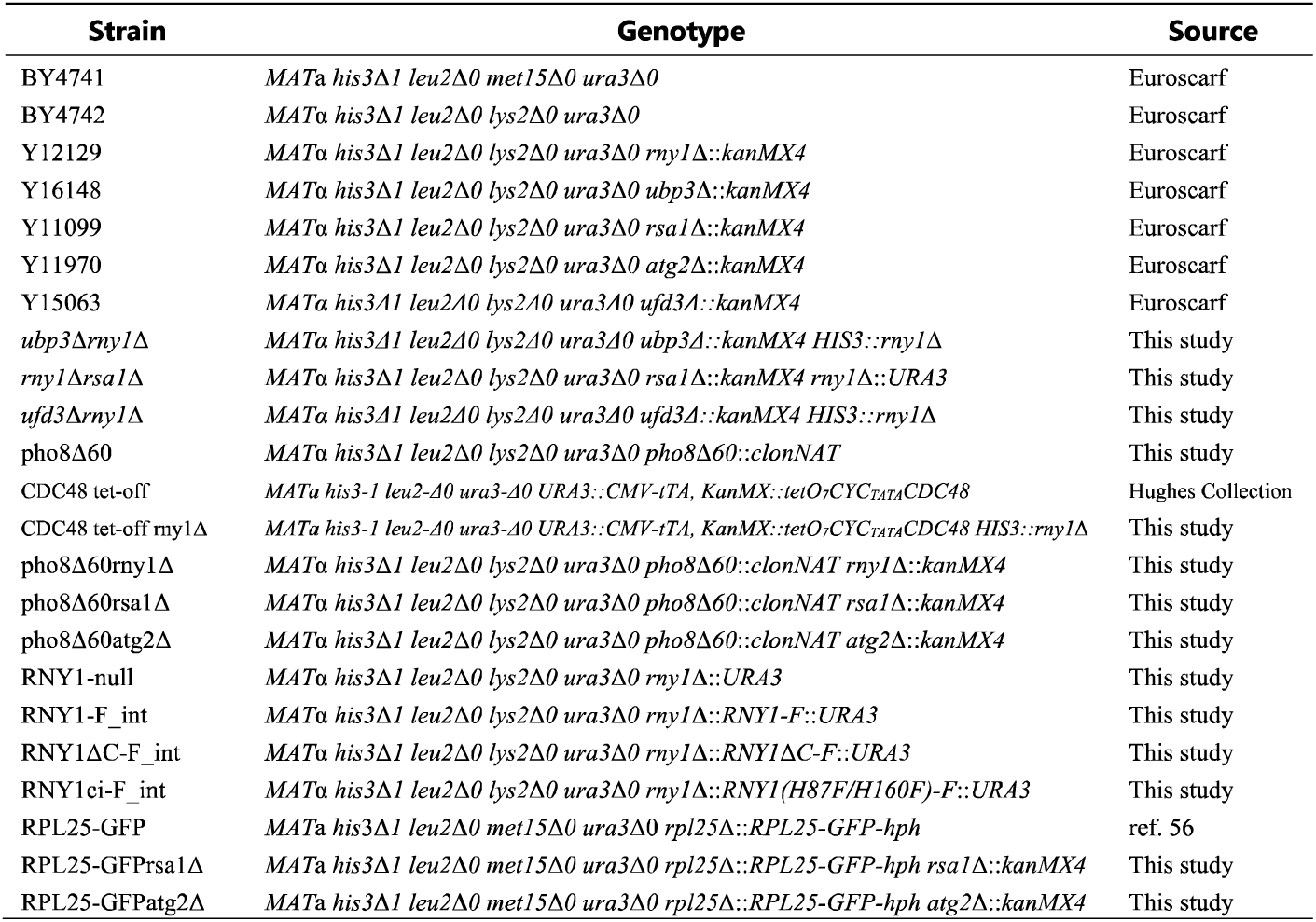
Yeast Strains used in this study.

**Table S1.** Top 5 hits for protein structures structurally similar to the C-terminal extension of Rny1p by Dali search.

| Rank | PDB ID | Z score | RMSD [Å] | Description |
| --- | --- | --- | --- | --- |
| 1 | 5B2H_A | 11.0 | 3.1 | HA-33 |
| 2 | 3AH2_B | 10.8 | 3.3 | MAIN HEMAGGLUTININ COMPONENT |
| 3 | 2VSE_A | 10.7 | 3.2 | MOSQUITOCIDAL TOXIN |
| 4 | 4IY8_A | 10.5 | 3.0 | 30K PROTEIN 1 |
| 5 | 4LO0_A | 10.5 | 3.3 | HA-33 |

## REFERENCES

1. Pestov, D.G. and Shcherbik, N. (2012) Rapid cytoplasmic turnover of yeast ribosomes in response to rapamycin inhibition of TOR. Mol. Cell. Biol., 32, 2135–2144.

2. Kitahara, K. and Miyazaki, K. (2011) Specific inhibition of bacterial RNase T2 by helix 41 of 16S ribosomal RNA. Nat. Commun., 2, 549.

3. Haud, N., Kara, F., Diekmann, S., Henneke, M., Willer, J.R., Hillwig, M.S., Gregg, R.G., Macintosh, G.C., Gärtner, J., Alia, A., et al. (2011) rnaset2 mutant zebrafish model familial cystic leukoencephalopathy and reveal a role for RNase T2 in degrading ribosomal RNA. Proc. Natl. Acad. Sci. U. S. A., 108, 1099–1103.

4. Hillwig, M.S., Contento, A.L., Meyer, A., Ebany, D., Bassham, D.C. and Macintosh, G.C. (2011) RNS2, a conserved member of the RNase T2 family, is necessary for ribosomal RNA decay in plants. Proc. Natl. Acad. Sci. U. S. A., 108, 1093–1098.

5. Andersen, K.L. and Collins, K. (2012) Several RNase T2 enzymes function in induced tRNA and rRNA turnover in the ciliate Tetrahymena. Mol. Biol. Cell, 23, 36–44.

6. Acquati, F., Morelli, C., Cinquetti, R., Bianchi, M.G., Porrini, D., Varesco, L., Gismondi, V., Rocchetti, R., Talevi, S., Possati, L., et al. (2001) Cloning and characterization of a senescence inducing and class II tumor suppressor gene in ovarian carcinoma at chromosome region 6q27. Oncogene, 20, 980–988.

7. 7. Henneke, M., Diekmann, S., Ohlenbusch, A., Kaiser, J., Engelbrecht, V., Kohlschütter, A., Krätzner, R., Madruga-Garrido, M., Mayer, M., Opitz, L., et al. (2009) RNASET2-deficient cystic leukoencephalopathy resembles congenital cytomegalovirus brain infection. Nat. Genet., 41, 773–775.

8. Ostendorf, T., Zillinger, T., Andryka, K., Schlee-Guimaraes, T.M., Schmitz, S., Marx, S., Bayrak, K., Linke, R., Salgert, S., Wegner, J., et al. (2020) Immune Sensing of Synthetic, Bacterial, and Protozoan RNA by Toll-like Receptor 8 Requires Coordinated Processing by RNase T2 and RNase 2. Immunity, 52, 591–605.e6.

9. Greulich, W., Wagner, M., Gaidt, M.M., Stafford, C., Cheng, Y., Linder, A., Carell, T. and Hornung, V. (2019) TLR8 Is a Sensor of RNase T2 Degradation Products. Cell, 179, 1264–1275.e13.

10. Liu, K., Sato, R., Shibata, T., Hiranuma, R., Reuter, T., Fukui, R., Zhang, Y., Ichinohe, T., Ozawa, M., Yoshida, N., et al. (2021) Skewed endosomal RNA responses from TLR7 to TLR3 in RNase T2-deficient macrophages. Int. Immunol., 33, 479–490.

11. Miyake, K., Shibata, T., Fukui, R., Sato, R., Saitoh, S.-I. and Murakami, Y. (2022) Nucleic Acid Sensing by Toll-Like Receptors in the Endosomal Compartment. Front. Immunol., 13, 941931.

12. Takayama, S. and Isogai, A. (2005) Self-incompatibility in plants. Annu. Rev. Plant Biol., 56, 467–489.

13. Hua, Z.-H., Fields, A. and Kao, T.-H. (2008) Biochemical models for S-RNase-based self-incompatibility. Mol. Plant, 1, 575–585.

14. Schneider, R., Unger, G., Stark, R., Schneider-Scherzer, E. and Thiel, H.-J. (1993) Identification of a Structural Glycoprotein of an RNA Virus as a Ribonuclease. Science, 261, 1169–1171.

15. Langedijk, J.P.M. (2002) Translocation activity of C-terminal domain of pestivirus Erns and ribotoxin L3 loop. J. Biol. Chem., 277, 5308–5314.

16. Makarov, A.A. and Ilinskaya, O.N. (2003) Cytotoxic ribonucleases: molecular weapons and their targets. FEBS Lett., 540, 15–20.

17. MacIntosh, G.C., Bariola, P.A., Newbigin, E. and Green, P.J. (2001) Characterization of Rny1, the Saccharomyces cerevisiae member of the T2 RNase family of RNases: unexpected functions for ancient enzymes? Proc. Natl. Acad. Sci. U. S. A., 98, 1018–1023.

18. Shcherbik, N. (2013) Golgi-mediated glycosylation determines residency of the T2 RNase Rny1p in Saccharomyces cerevisiae. Traffic, 14, 1209–1227.

19. Kraft, C., Deplazes, A., Sohrmann, M. and Peter, M. (2008) Mature ribosomes are selectively degraded upon starvation by an autophagy pathway requiring the Ubp3p/Bre5p ubiquitin protease. Nat. Cell Biol., 10, 602–610.

20. Tatehashi, Y., Watanabe, D. and Takagi, H. (2016) γ-Glutamyl kinase is involved in selective autophagy of ribosomes in Saccharomyces cerevisiae. FEBS Lett., 590, 2906–2914.

21. Zhang, T., Lei, J., Yang, H., Xu, K., Wang, R. and Zhang, Z. (2011) An improved method for whole protein extraction from yeast Saccharomyces cerevisiae. Yeast, 28, 795–798.

22. Noda, T. and Klionsky, D.J. (2008) The quantitative Pho8Delta60 assay of nonspecific autophagy. Methods Enzymol., 451, 33–42.

23. Love, J., Selker, R., Marsman, M., Jamil, T., Dropmann, D., Verhagen, J., Ly, A., Gronau, Q.F., Šmíra, M., Epskamp, S., et al. (2019) JASP: Graphical Statistical Software for Common Statistical Designs. J. Stat. Softw., 88, 1–17.

24. Jumper, J., Evans, R., Pritzel, A., Green, T., Figurnov, M., Ronneberger, O., Tunyasuvunakool, K., Bates, R., Žídek, A., Potapenko, A., et al. (2021) Highly accurate protein structure prediction with AlphaFold. Nature, 596, 583–589.

25. Mirdita, M., Schütze, K., Moriwaki, Y., Heo, L., Ovchinnikov, S. and Steinegger, M. (2022) ColabFold: making protein folding accessible to all. Nat. Methods, 19, 679–682.

26. ColabFold - Making protein folding accessible to all, Protein complex prediction with AlphaFold-Multimer.

27. Li, W. and Godzik, A. (2006) Cd-hit: a fast program for clustering and comparing large sets of protein or nucleotide sequences. Bioinformatics, 22, 1658–1659.

28. Matsui, M. and Iwasaki, W. (2020) Graph Splitting: A Graph-Based Approach for Superfamily-Scale Phylogenetic Tree Reconstruction. Syst. Biol., 69, 265–279.

29. Huang, H., Kawamata, T., Horie, T., Tsugawa, H., Nakayama, Y., Ohsumi, Y. and Fukusaki, E. (2015) Bulk RNA degradation by nitrogen starvation-induced autophagy in yeast. EMBO J., 34, 154–168.

30. Ossareh-Nazari, B., Bonizec, M., Cohen, M., Dokudovskaya, S., Delalande, F., Schaeffer, C., Van Dorsselaer, A. and Dargemont, C. (2010) Cdc48 and Ufd3, new partners of the ubiquitin protease Ubp3, are required for ribophagy. EMBO Rep., 11, 548–554.

31. Wyant, G.A., Abu-Remaileh, M., Frenkel, E.M., Laqtom, N.N., Dharamdasani, V., Lewis, C.A., Chan, S.H., Heinze, I., Ori, A. and Sabatini, D.M. (2018) NUFIP1 is a ribosome receptor for starvation-induced ribophagy. Science, 360, 751–758.

32. McKeegan, K.S., Debieux, C.M., Boulon, S., Bertrand, E. and Watkins, N.J. (2007) A dynamic scaffold of pre-snoRNP factors facilitates human box C/D snoRNP assembly. Mol. Cell. Biol., 27, 6782–6793.

33. Boulon, S., Marmier-Gourrier, N., Pradet-Balade, B., Wurth, L., Verheggen, C., Jády, B.E., Rothé, B., Pescia, C., Robert, M.-C., Kiss, T., et al. (2008) The Hsp90 chaperone controls the biogenesis of L7Ae RNPs through conserved machinery. J. Cell Biol., 180, 579–595.

34. Quinternet, M., Rothé, B., Barbier, M., Bobo, C., Saliou, J.-M., Jacquemin, C., Back, R., Chagot, M.-E., Cianférani, S., Meyer, P., et al. (2015) Structure/Function Analysis of Protein–Protein Interactions Developed by the Yeast Pih1 Platform Protein and Its Partners in Box C/D snoRNP Assembly. J. Mol. Biol., 427, 2816–2839.

35. Rothé, B., Back, R., Quinternet, M., Bizarro, J., Robert, M.-C., Blaud, M., Romier, C., Manival, X., Charpentier, B., Bertrand, E., et al. (2014) Characterization of the interaction between protein Snu13p/15.5K and the Rsa1p/NUFIP factor and demonstration of its functional importance for snoRNP assembly. Nucleic Acids Res., 42, 2015–2036.

36. Birgisdottir, Å.B., Lamark, T. and Johansen, T. (2013) The LIR motif - crucial for selective autophagy. J. Cell Sci., 126, 3237–3247.

37. Evans, R., O’Neill, M., Pritzel, A., Antropova, N., Senior, A., Green, T., Žídek, A., Bates, R., Blackwell, S., Yim, J., et al. (2021) Protein complex prediction with AlphaFold-Multimer. bioRxiv, 10.1101/2021.10.04.463034.

38. Floyd, B.E., Morriss, S.C., MacIntosh, G.C. and Bassham, D.C. (2015) Evidence for autophagy-dependent pathways of rRNA turnover in Arabidopsis. Autophagy, 11, 2199–2212.

39. Holm, L. (2020) DALI and the persistence of protein shape. Protein Sci., 29, 128–140.

40. Sagane, Y., Hayashi, S., Akiyama, T. and Matsumoto, T. (2016) Conformational divergence in the HA-33/HA-17 trimer of serotype C and D botulinum toxin complex. Biochemical and.

41. Kobayashi, H., Itagaki, T., Inokuchi, N., Ohgi, K., Wada, T., Iwama, M. and Irie, M. (2003) A new type of RNase T2 ribonuclease in two Basidiomycetes fungi, Lentinus edodes and Irpex lacteus. Biosci. Biotechnol. Biochem., 67, 2307–2310.

42. Hulst, M.M., Himes, G., Newbigin, E. and Moormann, R.J. (1994) Glycoprotein E2 of classical swine fever virus: expression in insect cells and identification as a ribonuclease. Virology, 200, 558–565.

43. Luhtala, N. and Parker, R. (2010) T2 Family ribonucleases: ancient enzymes with diverse roles. Trends Biochem. Sci., 35, 253–259.

44. Floyd, B.E., Mugume, Y., Morriss, S.C., MacIntosh, G.C. and Bassham, D.C. (2017) Localization of RNS2 ribonuclease to the vacuole is required for its role in cellular homeostasis. Planta, 245, 779–792.

45. Liu, Y., Zou, W., Yang, P., Wang, L., Ma, Y., Zhang, H. and Wang, X. (2018) Autophagy-dependent ribosomal RNA degradation is essential for maintaining nucleotide homeostasis during C. elegans development. Elife, 7.

46. Bardoni, B., Willemsen, R., Weiler, I.J., Schenck, A., Severijnen, L.-A., Hindelang, C., Lalli, E. and Mandel, J.-L. (2003) NUFIP1 (nuclear FMRP interacting protein 1) is a nucleocytoplasmic shuttling protein associated with active synaptoneurosomes. Exp. Cell Res., 289, 95–107.

47. Rothé, B., Saliou, J.-M., Quinternet, M., Back, R., Tiotiu, D., Jacquemin, C., Loegler, C., Schlotter, F., Peña, V., Eckert, K., et al. (2014) Protein Hit1, a novel box C/D snoRNP assembly factor, controls cellular concentration of the scaffolding protein Rsa1 by direct interaction. Nucleic Acids Res., 42, 10731–10747.

48. Makino, S., Kawamata, T., Iwasaki, S. and Ohsumi, Y. (2021) Selectivity of mRNA degradation by autophagy in yeast. Nat. Commun., 12, 2316.

49. Samir, P., Browne, C.M., Rahul, Sun, M., Shen, B., Li, W., Frank, J. and Link, A.J. (2018) Identification of changing ribosome protein compositions using mass spectrometry. Proteomics, 18, e1800217.

50. Callaway, E. (2022) What’s next for AlphaFold and the AI protein-folding revolution. Nature, 604, 234–238.

51. Humphreys, I.R., Pei, J., Baek, M., Krishnakumar, A., Anishchenko, I., Ovchinnikov, S., Zhang, J., Ness, T.J., Banjade, S., Bagde, S.R., et al. (2021) Computed structures of core eukaryotic protein complexes. Science, 374, eabm4805.

52. Khvorova, A., Kwak, Y.G., Tamkun, M., Majerfeld, I. and Yarus, M. (1999) RNAs that bind and change the permeability of phospholipid membranes. Proc. Natl. Acad. Sci. U. S. A., 96, 10649–10654.

